# A piece-wise linear recurrent neural network identifies generalizable dynamics from neural activity during decision-making in rodents

**DOI:** 10.64898/2026.09.29.754775

**Authors:** William H. Barnett, Daniel Durstewitz, Alexey Kuznetsov, Christopher C. Lapish

## Abstract

Alterations in impulsivity characterize several neuropsychiatric disorders, however the neural processes that underlie an impulsive choice have not been identified. A recent body of literature suggests that there may be generalizable aspects of neural representations across animals. Whether this is also true for dynamic rather than just static (like geometrical) properties of neural representations, and specifically in the context of impulsive behaviors, is less clear. Therefore, the goal of this study is to characterize *generalizable* features of latent dynamics that disambiguate an impulsive from non-impulsive choice across many different data sets and animals. A deep learning-based latent factor model was constructed from neural recordings that were acquired in rat anterior cingulate cortex while performing a delay discounting task, which is commonly used to assess impulsivity. Prior work shows this brain region is critical for the performance of this task. We leveraged piecewise linear recurrent neural networks (PLRNNs) to perform a nonlinear latent factor analysis across animals and experimental sessions. Our PLRNN explained a substantially larger percent of variance in fewer dimensions than linear approaches. We cross-validated performance of the model on test data and following the disruption of recurrent connectivity or auto-regressive latent dynamics. These tests confirmed that our fitting procedure consistently embedded generalizable neuronal dynamics within the PLRNN. Further, we found that the latent dynamics exhibited strong rotational features and progressed faster for impulsive choices. In summary, our model captures properties of the latent dynamics that are consistent across multiple data sets and disambiguate impulsive choices.

## Introduction

The anterior cingulate cortex (ACC) plays a critical role in several aspects of decision making, including intertemporal choices (Floresco et al., 1997; Gisquet-Verrier & Delatour, 2006; Goto & Grace, 2008; Myroshnychenko et al., 2017; van Heukelum et al., 2020). The ACC is highly connected to many other brain regions, and it is also highly recurrent (Carr & Sesack, 2000; Hoover & Vertes, 2007; Lévesque & Parent, 1998; van Heukelum et al., 2020). Due to this connectivity, the ACC is positioned to link highly processed sensory input to the implementation of actions (Euston et al., 2012). As a nexus for cognitive control of behavior, the ACC is also implicated in dysfunctional decision making including the lack of impulsive control (Churchwell et al., 2009; Dalley et al., 2011; Loos et al., 2010; McLaughlin & Redish, 2023), and recently, we have shown how inactivation of ACC promotes impulsivity, the readout of which is an increase in immediate choices (White et al., 2023). To understand the mechanisms that underlie an intertemporal choice, it is necessary to investigate temporal evolution of neural activity (i.e. neural dynamics) during decision making in the ACC.

A foundational model of perceptual decision making attributes the mechanism of choice to the activity of choice-specific neurons that integrate information until a threshold for action is reached (Wong & Wang, 2006). This approach conceptualizes a choice as evolving in a decision space where neural activity accumulates evidence available from sensory inputs in order to perform one choice or the other (Brody & Hanks, 2016; Gold & Shadlen, 2007; Hanks & Summerfield, 2017; Kurikawa et al., 2018).

Dimensionality reduction approaches have proven useful to distill a lower-dimensional space from large groups of neurons that captures the principal features of neural dynamics. Task-related activity is thought to occupy relatively low-dimensional manifolds within a high-dimensional neural activity space (Gallego et al., 2017), and the neural activation associated with specific choices have been proposed to correspond to attractor states (Miller, 2016; Rabinovich & Varona, 2011). Most approaches to assess attractor-like dynamics in neuroscience so far mainly depend on visual inspection or statistical assessment of convergence properties of trajectories in neural state space projections obtained by dimensionality reduction techniques (Cunningham & Yu, 2014; Gallego et al., 2017; Safaie et al., 2023; Timme et al., 2022; White et al., 2023). Signatures of attractor-like dynamics have been detected in neural recordings from the ACC in the form of neural trajectories dwelling in regions of state space associated with task-related behavior such as a specific choice (Balaguer-Ballester et al., 2011; Lapish et al., 2008, 2015). However, the aforementioned techniques lack a formal mathematical model that would describe the temporal evolution of trajectories in state space as a dynamical system. Further, a formal model allows for predictions to be made about how latent variables might interact to facilitate intertemporal choices.

Data driven models of neural population activity are attractive as a dimensionality reduction tool and for their interpretability (Macke et al., 2011; Nonnenmacher et al., 2017; Sussillo et al., 2015; Zhao & Park, 2016). Machine learning models based on recurrent neural networks (RNNs) have been developed to investigate how neural activity may generate animal behavior across individual trials (Durstewitz, 2017; Pandarinath et al., 2018; Koppe et al., 2019; Keshtkaran et al., 2022; Sedler & Pandarinath, 2023; Patel et al., 2023) and across multiple regions of the brain (Perich et al., 2021). This approach may be further leveraged by using RNNs with low rank connectivity to recover low-dimensional latent dynamics from neural populations in which latent dynamics are constrained to a sub-space of the latent space (Valente et al., 2022; Hess et al., 2023; Brenner, Hess, et al., 2024; Pals et al., 2024). We contribute to this body of work by investigating generalization of neural latent states across multiple animals performing a decision-making task.

Generally, RNNs are universal approximators of dynamical systems (Funahashi & Nakamura, 1993; Kimura & Nakano, 1998; Chow & Li, 2000). Piecewise linear RNNs (PLRNNs) are a class of RNNs that combine tractability due to piecewise linear activation with the utility of nonlinear models; moreover, they are capable of generating accurate neural activity patterns when inferred directly from data sets measured from the brain (Durstewitz, 2017; Durstewitz et al., 2023; Koppe et al., 2019). Here, we leverage a PLRNN as a data-driven model to investigate latent brain states and recover relatively low-dimensional dynamics. Currently, the extent to which inferred low-dimensional neural dynamics are capable of generalizing to multiple different and independent subjects and datasets is an open question (Melbaum et al., 2022; Safaie et al., 2023). To attack this problem, we address the latent neural factors that underlie intertemporal choice and how they are altered in impulsive decisions. We used an adjusting amount delay discounting task designed to assess impulsivity in rats (Linsenbardt et al., 2017; Seamans et al., 2024; White et al., 2023). Specifically, we test the hypothesis that a single PLRNN model can capture the neural dynamics of animals performing an intertemporal choice and whether the latent dynamical factors provided by such a model can generalize across datasets collected from different animals. If successful, this will be a powerful tool to better understand generic computational mechanisms underlying brain functions and their impairments.

## Results

### Description of Data

In vivo neural recordings were acquired from the anterior cingulate cortex of awake behaving rats performing an adjusting amount delay discounting task. The critical part of this task is where an animal must press a lever to choose between an immediate, small reward or a large, delayed reward, and impulsivity is characterized as a greater propensity to choose the immediate lever. Neuronal recordings were grouped by choice (immediate and delay), synchronized to time of the lever press that reports the choice and averaged across trials within each session. Spike trains were binned at 200ms and smoothed with an adaptive Gaussian filter that was inversely proportional to the mean interspike interval (ISI): σ=1/ISI. After averaging, we selected neurons that exhibited non-zero firing rates for both immediate and delay choices. Following this preprocessing, we selected experimental recordings with at least 5 neurons; this selection resulted in 52 recordings from 9 animals containing in total 2563 neurons.

### PLRNN accurately reproduces neuronal dynamics of decision making

We trained PLRNNs to reproduce trial-averaged neuronal activity recorded in rats during the delay discounting task (Fig. 1A). Our model was configured to generate the latent dynamics required to reproduce trial-averaged neural activity for either an immediate or delay choice depending on the initial condition, which was a trained model parameter (Fig. 1B). The total pool of 52 recordings were randomly partitioned into a training dataset that contained 10 recordings and a test dataset that contained 42 recordings. The training of the model produced a single latent model that described the trial-averaged neuronal activity for both choice groups (Fig. 1C).

**Figure 1.**
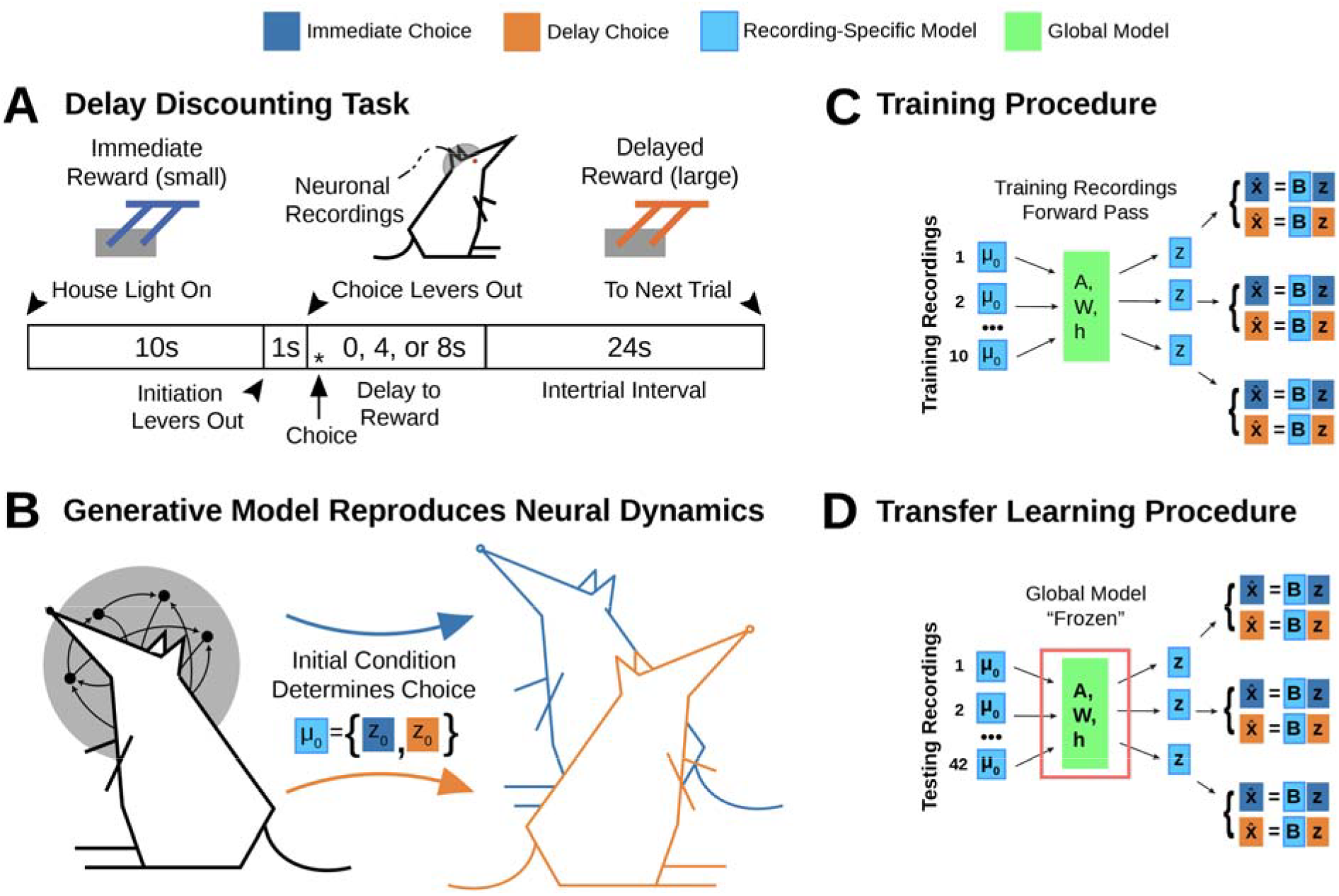
Methods for constructing a PLRNN to reproduce neuronal dynamics of anterior cingulate cortex during a delay discounting task. (A) We acquired multiple single-unit recordings from awake behaving animals during a delay discounting task. This data is synchronized to the time of the choice, transformed to 200 ms time-binned histograms, and trial averaged by choice. In one model instance, 10 of 52 recordings were randomly selected for use as the training dataset, and the remaining 42 were used for testing. (B) Training and testing recordings contained the trial averaged neuronal firing rates for separate immediate choice and delay choice activity. As such, certain model quantities were specialized to choice group: the latent model was trained to reflect both choices, and the choice was encoded by the initial condition ***μ***_*0*_. (C) Optimization of the latent model was performed on the 10 recordings of neuronal activity in the training dataset. Since the initial condition ***μ***_*0*_ and observation model *B* for each recording were specific to the neuronal activity within that recording, the parameters ***μ***_*0*_ and B were fit separately for each recording in the training dataset. The parameters ***A, W***, and ***h*** were responsible for producing shared dynamics and were shared across all training recordings. Since each recording produced a set of predicted observations for both immediate and delay choices, we averaged mean absolute errors over time, over neurons, and over choice. (D) Our PLRNN was evaluated against recordings of neuronal activity from the testing dataset. In order to produce performance metrics, it was necessary to fit the recording-specific values for the initial condition of the latent model and for the observation model. These parameters ***μ***_*0*_ and ***B*** were fit much the same way as the model was trained in the training dataset, except the latent model parameters were “frozen” (held fixed) while the recording-specific initial condition and observation model parameters were fit.

When fitting the parameters of the latent model (***A, W***, and ***h***), it was necessary to also fit the initial condition (***μ***_*0*_) and observation model (***B***) for each recording, since each recording consists of different units with different mappings from the latent space to the observations. Thus, it was necessary to fit new values for the parameters ***μ***_*0*_ and ***B*** for each recording in the testing dataset. During this operation, the latent model parameters ***A, W***, and ***h*** were “frozen” such that they were not further optimized (Fig 1D). Since it was necessary to perform separate fitting of the observation model for the testing dataset, we incorporated L1 and L2 penalties in fitting observation parameters ***B*** to reduce the risk of overfitting (Fig S1).

Within a model instance, generated latent dynamics (Fig 2A) were used to produce predicted firing rates for each neuron (Fig 2B). The fitting of model instances was accomplished by minimizing mean absolute error (MAE) between observed and predicted firing rates of each neuron. For the short trial-averaged time series of 151 data points, we evaluated model fits by MAE (Fig 2C) as well as by computing the Pearson correlation coefficient between observed and predicted firing rates (Fig 2D). To evaluate model fits for all individual neurons, the Pearson correlation coefficients were averaged across the immediate and delay trials (AKA choice groups). These results indicate an accurate reproduction of trial-averaged neuronal data by our PLRNN.

**Figure 2.**
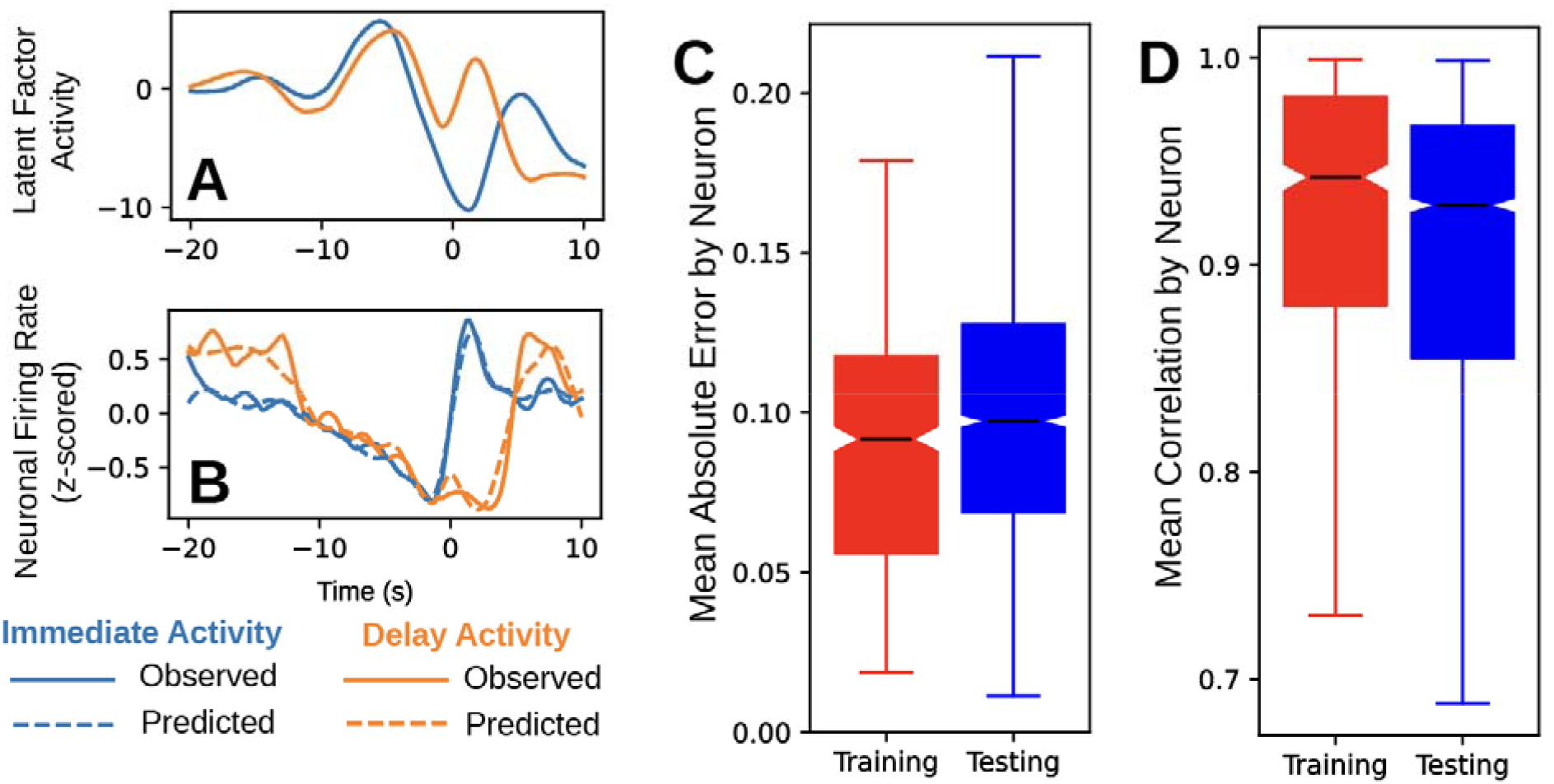
PLRNN fitting procedure reproduces neuronal dynamics of delay discounting. Model simulations produce distinct immediate and delayed choice latent model solutions (A) and closely reproduce the firing rates recorded in both choices (B). The performance of the model in one model instance was quantified by mean absolute error (C) and Pearson’s correlation coefficient (D) with 10 neuronal recordings in the training dataset and 42 neuronal recordings in the testing dataset.

### Dynamics of the latent model are critical to accurately generate firing rates

Our fitting routine trained a single latent model that generalized across 52 recordings from 9 animals. These results indicated that generic properties of the trial-averaged neuronal activity profiles associated with decision making were extracted from the set of training data and embedded as structure in the weights of our model. However, it was not clear whether (1) the decision dynamics were captured by the latent model and then loaded onto individual neurons by the observation model, or (2) the latent model acted as merely a reservoir of sufficient complexity that could be loaded onto the neuronal activity by the observation model, i.e. a form of overfitting induced during the fitting of the observation factor loading matrix ***B***. To distinguish between these two scenarios, we performed two numerical experiments to control for overfitting and cross validated these experiments over 100 independent model instances.

To facilitate overfitting experiments, we introduced a hyperparameter that determined the size of the latent model (*M*) to illustrate cases (1) and (2). The number of latent factors available for producing the dynamics is the primary hyperparameter of our RNN, and model selection required the investigation of the size of the latent model as well as several experiments to evaluate overfitting in the observation model. To evaluate the role of *M* in our PLRNN fit, we produced many (100) model instances for a broad range of values for *M* (Fig. 3A1). For each model instance, we independently sampled 10 recordings into the training dataset and the remaining 42 recordings into testing datasets.

**Figure 3.**
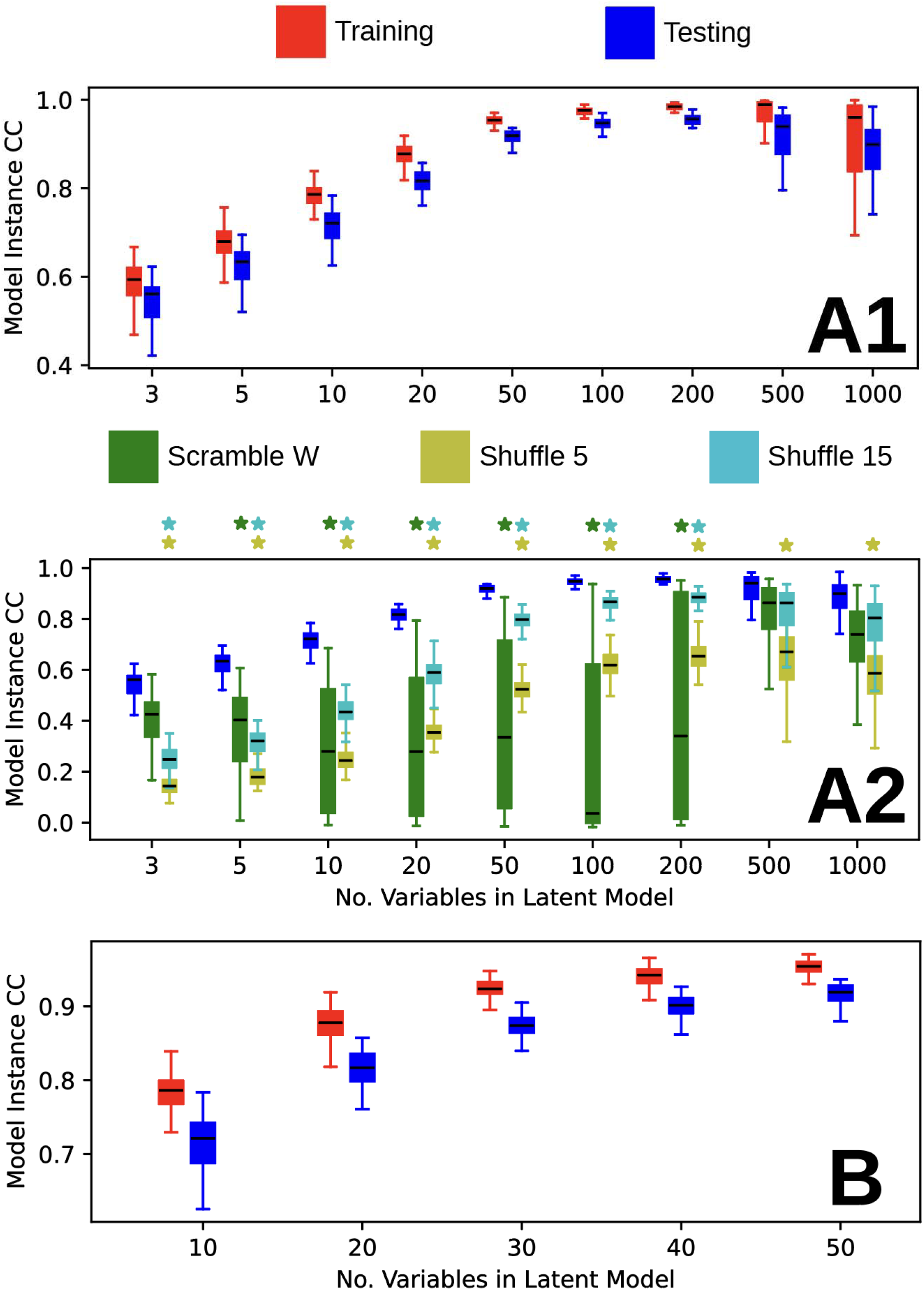
Optimal model performance was determined by the size of the latent model. Model instance correlation coefficient (CC) was determined by the correlation coefficient of the generated and observed spike trains and then averaged across neurons and across immediate and delay choices to produce a single value for each model instance. (A1) We depict the results from the training and testing datasets for a broad range of model sizes. (A2) In the group Scramble W, the off-diagonal elements of the **W** parameter were randomly scrambled, and the initial conditions and observation model were fit again for testing recordings. In the groups Shuffle 5 and Shuffle 15, the latent model time series for each testing recording was broken up into chunks of 5 and 15 time steps. These chunks were shuffled, and a new observation model was fit to these shuffled latent model solutions. We depict the results of the control groups Scramble W, Shuffle 5, and Shuffle 15 in comparison to the testing group. Each of the perturbations of the latent model or latent model activity degraded the fits of the observations therefore highlighting the importance of the latent dynamics. Stars above each model size indicate statistical significance of a bootstrap test for overlap of the IQR between the testing performance distribution and the Scramble-W, Shuffle 5, or Shuffle 15 performance distributions. These tests were performed with 10000 bootstrap samples each. The color code indicates which test is performed; for example the green star where the model size is 10 indicates a significant test of the Scramble W group against the testing group. Here, the alpha level was adjusted with the Bonferroni correction for multiple comparisons considering 27 comparisons (alpha = 0.05 / 27, which is approximately 1.85e-3). (B) We inspected model sizes from 10 to 50 in increments of to select the size of the reported model. The training and testing distributions are depicted.

Here we report the performance for groups of individual model instances by averaging neuron Pearson correlation coefficients across choice group and then over all neurons in the set (i.e. the training dataset). We observed that training and test dataset performance increased with the size of the latent model up to approximately *M*=200 (Fig. 3A1). For larger values of *M*, the median test performance dropped. We next used these test performance results for comparison in two overfitting experiments.

In the first overfitting experiment, we disrupted the recurrent dynamics of the latent model by scrambling the off-diagonal weights in the recurrent connectivity (the matrix ***W***), termed ‘Scramble-W’ in the following, and then refit the initial condition and observation model for the testing recordings (following the procedure described for the test dataset) (Fig. 3A2). The model performance in the Scramble-W experiments varied greatly. We quantified the performance of models in this experiment with a bootstrap test for overlapping interquartile range (IQR). The Scramble-W performance distributions were significantly lower than the test model performance distributions for values of *M* from 5 to 200 (Fig 3A2). Outside of this range, the Scramble-W performance was similar to that of the test performance. These results indicate overfitting of the observation model for relatively large values of the hyperparameter *M*, indicating the importance of latent model dynamics for reproducing the observations.

In the second overfitting experiment, we disrupted the dependence of the latent model on its history by shuffling the time series of the unperturbed latent model in chunks of 5 or 15 time steps before refitting the observation model. These results were named Shuffle 5 and Shuffle 15 (Fig 3A2). We quantified the performance of the Shuffle 5 and Shuffle 15 experiments with a bootstrap test for overlapping IQR. For all values of the hyperparameter *M*, the Shuffle 5 performance was significantly different compared to the testing performance. The performance for Shuffle 15 was significantly lower than testing performance for lower values of *M*. However, where *M* was 500 and 1000, the performance for Shuffle 15 became similar to the performance in the testing dataset. We interpret the Shuffle 15 results to indicate overfitting of the observation model for larger values of *M*, which is expected as the size of ***B*** grows as *M×N*, with *N* the number of observed units.

We integrated over this set of control experiments to inform model selection. For values of *M* from 20 to 50, our Scramble-W and Shuffle performance results exhibited the largest difference from that of the testing dataset (Fig 3A2), and we focused on this range to select a model for further evaluation (Fig 3B). In this range, model instances exhibited a trade-off between the size of the latent model and testing performance. At *M*=30 the testing performance was satisfactory, and there was clear separation between the performance of the PLRNN and our control experiments across model instances. These results indicate the successful embedding of neuronal dynamics within the latent model. In the model instances where *M*=30, we selected the instance with lowest MAE in the test dataset that is shown in Fig 2.

### Latent dynamics generate neural activity patterns in a consistent manner across model instances

If our fitting routine consistently reconstructed neural dynamics, then the features that generate neuronal activity should be constructed in similar ways across model instances. We assessed this prediction in a quantitative analysis of the observation model (the matrix ***B***). Neurons that exhibited similar activity should be loaded by the observation model onto the latent model in a similar fashion across model instances. To assess this, we used k-means clustering to group neurons by their loadings in ***B*** in each model instance (visual aid; Fig 4A). In one model instance, neurons within a single cluster have similar loading coefficients in the observation model compared to neurons in different clusters. If different neurons are consistently found within the same cluster over many model instances, then we conclude that they are loaded in the observation model in a similar fashion across model instances. To quantify these results, we counted the frequency with which each neuron shared a cluster with each other neuron over model instances and constructed a histogram of the values in this matrix (Fig 4B). Neurons from the same recording may bias this analysis as they may have greater correlations and, thus, may produce very similar loadings. To control for the possibility that the heavy tail of our distribution could be explained by neurons observed within the same recording, we excluded those counts and repeated this analysis (Fig 4C). This distribution was significantly different from null distributions that were generated by shuffling the cluster labels among neurons (sampled 100 null distributions; two-sample KS test: D(6382994)=0.149 p-values < 0.0001). These results indicate that our fitting routine loaded neurons in the observation model in a consistent manner across model instances.

**Figure 4.**
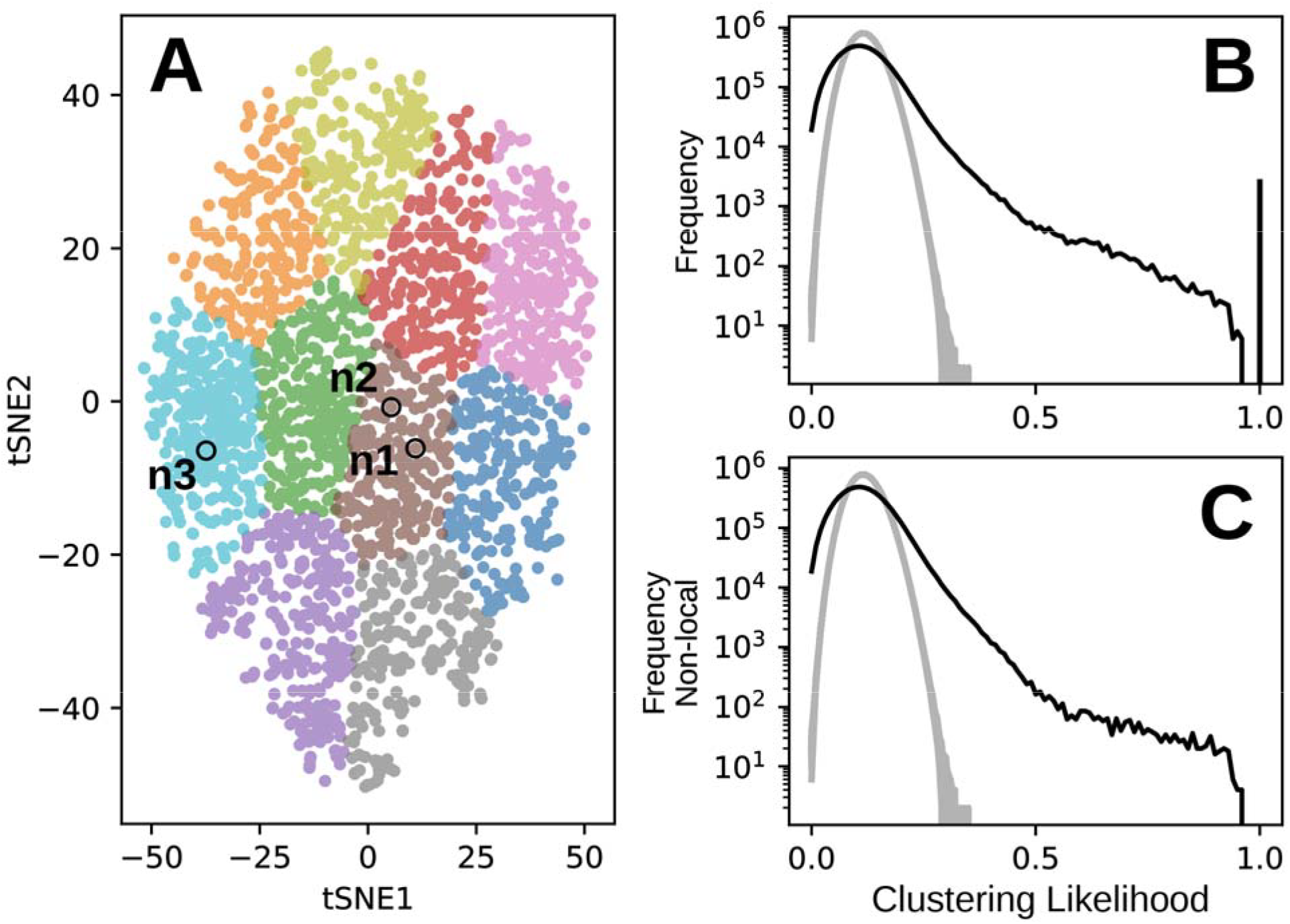
Latent model clustering likelihood distributions indicate stability of neuronal loadings in the observation model across model instances. (A) Colored points together represent a two-dimensional tSNE projection of the 30 features in the loading model (the matrix ***B***), and each point corresponds to the loading of an individual neuron onto the latent model. In this exemplar model instance, ten clusters are visualized by the color of the data. The neurons *n1* and *n2* are both found within the same cluster, and the neuron *n3* is in a different cluster. Since neuron *n2* is relatively close to neuron *n1* compared to neuron *n3*, neuron *n1* and *n2* are more likely belong to the same cluster in other model instances if our fitting routine consistently embeds features of neuronal activity into the observation model. (B) Black trace indicates the distribution of the probability of neuronal shared cluster participation over all model instances (N=100; description in **Methods**). The grey traces represent 100 computed null distributions produced by shuffling cluster labels. (C) This analysis was repeated after excluding entries in the clustering likelihood matrix that corresponded to interactions of neurons in the same recording.

### PLRNN fitting infers similar latent dynamics across model instances

From the above results we distilled the PLRNN with the minimal number of latent factors necessary to achieve a good reproduction of the neural dynamics. To evaluate the contribution of the nonlinear activation to embedding performance, we compared model to performance to two alternative linear approaches. We performed Principal Component Analysis (PCA) on the latent states of the PLRNN and compared these results to 1) PCA performed directly on the observed neural firing rates (Fig 5A) and 2) PCA performed on a version of our RNN with a linear activation function (Linear RNN abbreviated as LRNN; Fig S2-S3). Immediate and delay neural firing rates or immediate and delay latent model activity were concatenated in time and thus jointly investigated with PCA (Fig 5; Fig S3). Immediate choice and delay choice neural activity were separately projected onto the first two principal components (PCs; PC1 and PC2) and the explained variance was quantified. PC1 and PC2 revealed choice-specific features in the pattern of neural firing rates that looked superficially similar to the latent factors of the PLRNN (Fig 5A1-2, compare to Fig. 2A,B). However, the combined explained variance of the first two PCs was only 39% (Fig 5A3). To quantify the embedding performance of the LRNN and PLRNN models, we performed PCA on the latent factors from each recording of the testing dataset of all 100 model instances (Fig 5B1-4). In the PLRNN, the combined first two PCs accounted for 70% of explained variance (Fig 5B5). In comparison, the combined first two PCs of LRNN models only accounted for an average of 35% of explained variance (Fig S3B5). Thus, our PLRNN latent factor analysis explained more variance in fewer dimensions compared to two linear techniques (LRNN and PCA) used to capture latent dynamics indicating that the PLRNN recovers more features of the neural dynamics.

**Figure 5.**
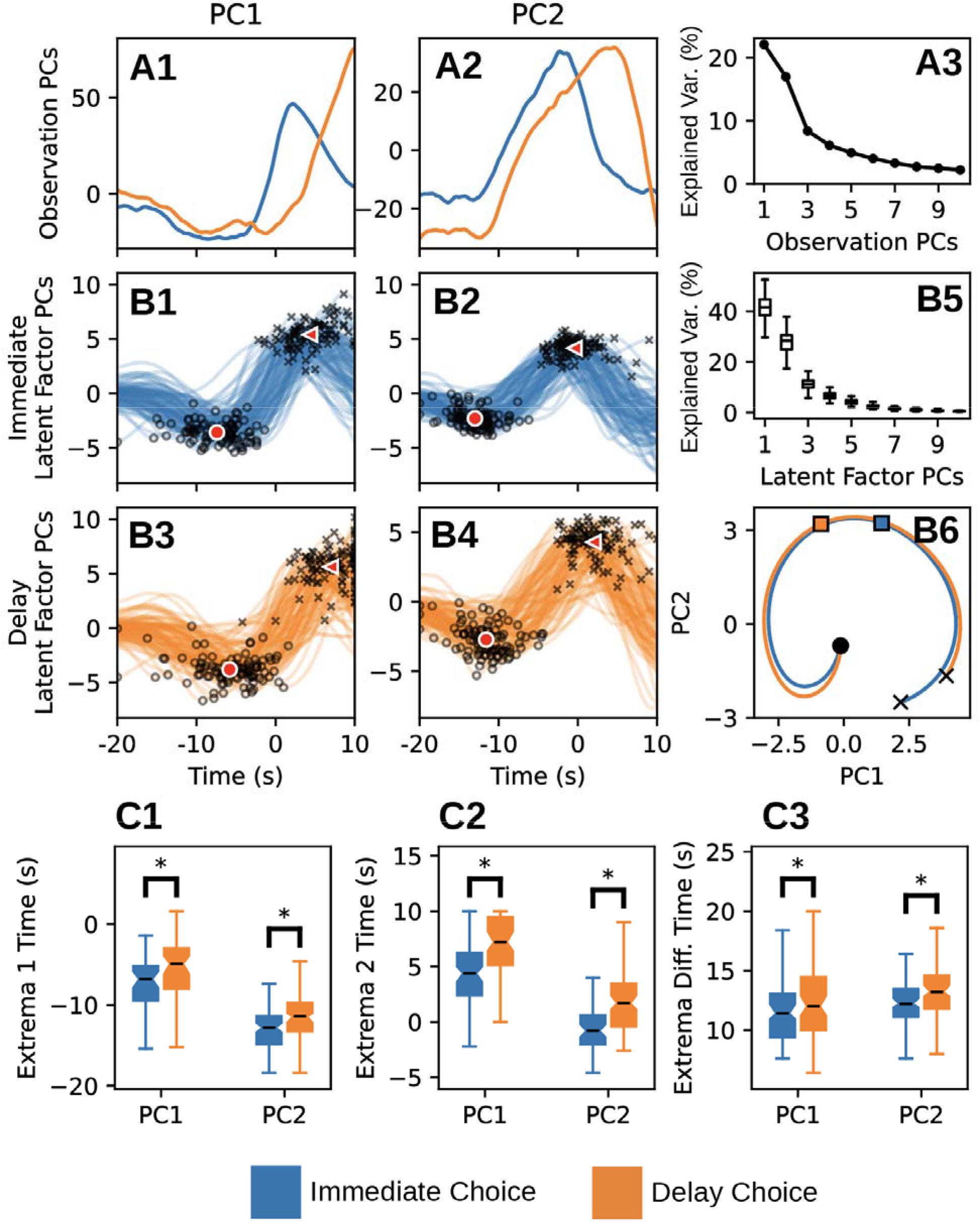
Latent model dynamics are consistent across PLRNN model instances and capture features of neuronal dynamics. In all panels, blue traces correspond to immediate choice activity, and orange traces correspond to delay choice activity. (A) Top row traces indicate results of PCA on neuronal activity depicting the PC1 (A1) and PC2 (A2) projections for immediate choice and delay choice. The combined explained variance of neuronal activity for the first two PCs was less than 50% (A3). PCA was performed separately on latent model activity for each model instance (B). Each colored line corresponds to the mean of the projections onto PC1 or PC2 for each of the 42 testing recordings within a model instance (B1-B4; one trace for each of 100 model instances). Latent dynamics in the PC1 and PC2 projections were characterized by the timing of two notable of extrema. The mean PC projection for each model instance is marked with a black circles to indicate Extrema 1 and a black X to indicate Extrema 2. The mean across model instances of the Extrema 1 positions is marked with a red dot, and the mean across model instances of the Extrema 2 positions is marked with a red triangle. The distribution of explained variance for the top 10 PCs for all model instances is visualized (B5). The mean of the traces from B1-B4 are adopted as generalized neuronal dynamics in the PC1-PC2 plot (B6). Within this projection, the progression of the state through time is marked: circles which correspond to the initial state; filled squares which indicate the time of the choice (time = 0); and Xs, which indicate the final state. Pairwise comparisons (independent t-test) for extrema timing across choice for PC1 and PC2 (C1-C3): * indicates p < 0.05. Extrema difference time is the time of Extrema 1 subtracted from the time of Extrema 2 (C3).

We sought to establish that we consistently reconstructed neuronal dynamics across model instances. Since our fitting routine assigned dynamical features onto latent factors in an unsupervised fashion, features of neuronal activity were distributed to latent factors in different ways from model instance to model instance. Simply put, from one model instance to the next, the activity pattern of individual latent factors could be very similar but exist in a different latent factor number. In a hypothetical example, reward-related features in one model instance might be captured by latent factors 2 and 3, but in a different model instance, these same features of neuronal activity could be captured by latent factors 4, 7, and 10. As such, it is difficult to directly compare the activity of latent factors across model instances. To identify the most prominent latent factor activity patterns, we utilized PCA. PCA was performed on latent model activity separately for each model instance. The first two principal components captured shared features of neuronal activity as it is embedded into the latent space (Fig 5B1-5B4). PCs were averaged across model instances and plotted in PC1-PC2 space; in this projection, latent model activity appeared to exhibit rotational dynamics (Fig 5B6).

The PC spaces recovered from the latent model appeared to exhibit rotational dynamics (Churchland et al., 2012); however, this analysis emphasized latent variables that were highly loaded onto the first 2 PCs and may not have captured generic properties of the un-altered latent model solutions. Therefore, we directly interrogated latent model solutions to determine if they possessed rotational characteristics. In the PLRNN, activation functions are zero for inputs below zero and linear for inputs above zero, and as such, the Jacobian of the latent model is constant within each linear subregion (see Methods). We computed the eigenvalues of the Jacobian for each linear subregion that was traversed by a latent model solution in each model instance (Fig 6A). Eigenvalues with imaginary components were observed in more than 99% of inspected subregions. However, trajectories in the latent space traversed many linear subregions: in half of model instances, trajectories traversed at least 49 unique subregions in 151 time steps (Fig 6B). These results indicated that the latent model possessed rotational components but were otherwise difficult to interpret since the eigenvalues abruptly changed as the latent state trajectory traversed multiple linear subregions. To capture the rotational properties of whole trajectories in the latent state space, we computed the gyration number—a model-free approach that is derived from the eigenvalues of the neuron differential covariance matrix (Kuzmina et al., 2024). The rotation strength is quantified by the imaginary component of the gyration number (referred to as the Rotation Axis in Fig 6C1). All model instances were above unity in the Decay-Rotation gyration plane (Fig 6C1), indicating a robust contribution of the rotational component to the dynamics. Moreover, the imaginary components of the gyration number for latent models were much larger than the imaginary component of the gyration number computed directly from the neural data (Fig 6C2). This indicates that latent model trajectories possess rotational dynamics, and these dynamics are enhanced in the latent model compared to the neural data.

**Figure 6.**
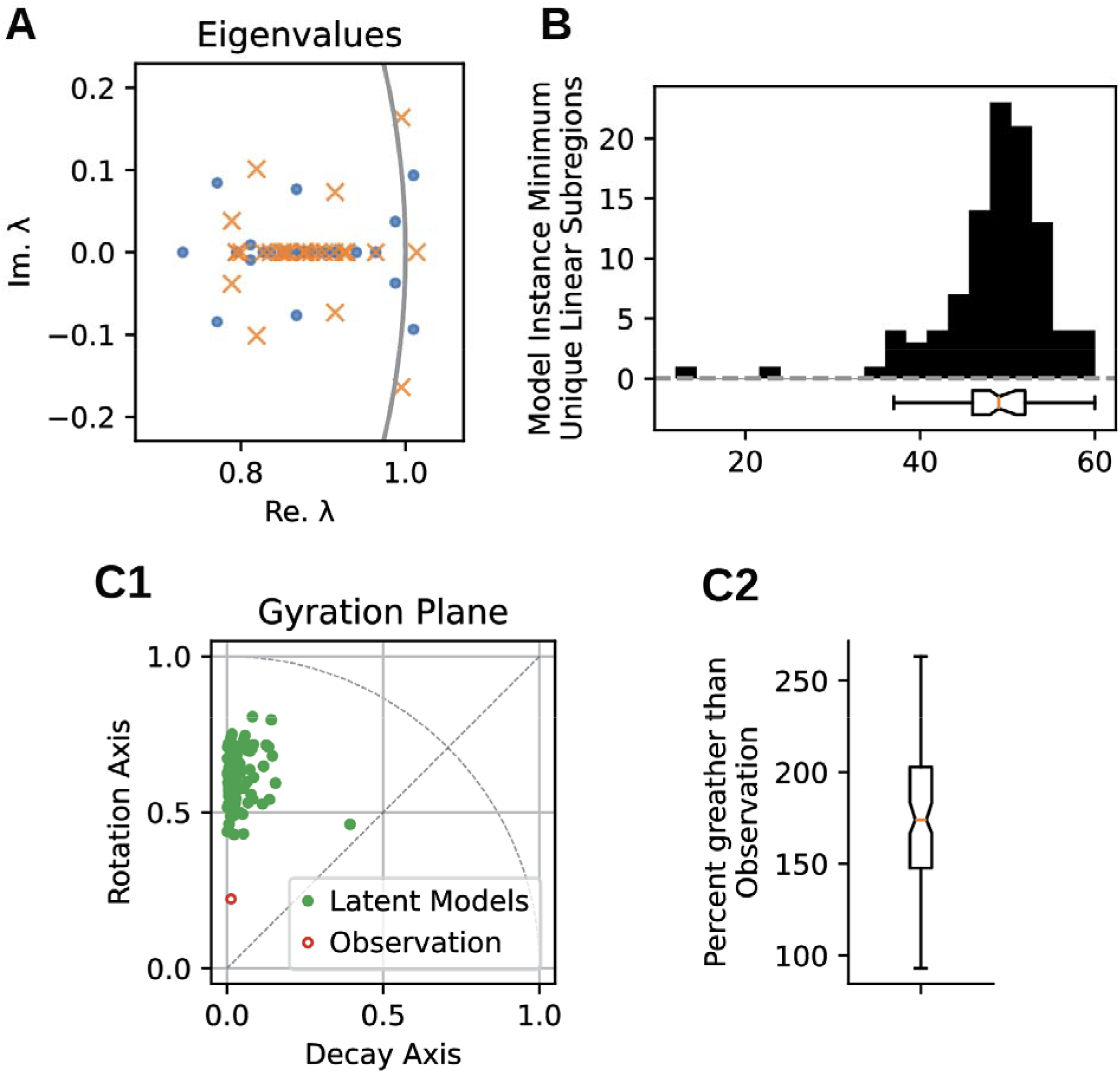
Characterization of rotational dynamics. (A) Eigenvalues of the Jacobian of the latent model at the time of the choice for immediate choice (blue) and delay choice (orange). (B) Each model instance contained 42 recordings in the testing group. For each session model instance, the number of traversed linear subregions were counted. We display here the distribution of the minimum such count for each model instance and quantify the same with a box-and-whisker plot. (C) Rotational dynamics were quantified using the gyration number. (C1) In the Gyration Plane, green data points correspond to the gyration number for the testing dataset from each model instance. The red point indicates the gyration number for the observed neural data. (C2) Box-and-whisker plot quantifies the distribution of the percent difference between the observed neural data and the group of latent model instances for the position of the gyration plane on the Rotation Axis.

The rotational phase of latent model activity appeared to be more advanced at the time of the choice in immediate vs delay choice (Fig 5B6). We quantified this difference in phase by characterizing prominent extrema in PC1 and PC2 projections of latent model activity across model instances (Fig 5C). PC1 and PC2 possessed an early extremum (Extrema 1) that usually preceded the time of the choice and a late extremum (Extrema 2) that was near or after the time of the choice (Fig 5B1-4). Extrema 1 occurred earlier in immediate choice for PC1 (two-sample unpaired t-test, t(198)=-3.0, p=2.7e-3) and PC2 (t(198)=-3.5, p=4.9e-4) compared to delay choice (Fig 5C1), and Extrema 2 occurred earlier in immediate choice for PC1 (t(198)=-6.8, t=9.9e-11) and PC2 (t(198)=-6.5, p=8.1e-10) compared to delay choice (Fig 5C2). We also examined the latency between Extrema 1 and Extrema 2 and found a shorter latency in immediate choice for PC1 (t(198)=-2.2, p=0.029) and PC2 (t(198)=-2.5, p=0.013) compared to delay choice (Fig 5C3). Taken together, these results indicate that features of neural activity occur earlier and progress faster in an immediate choice.

### Specialization of latent model dynamics to choice

Since the latent model differed only in initial conditions between immediate and delay choice simulations, the question remains how this gives rise to choice-specific activity profiles. We investigated specialization of the activity of latent factors in an exemplar model instance using the results of the aggregate PCAs described above; the computed neural dynamics of decision-making progressed in a rotational fashion from the initial conditions with differences in timing and extent between immediate and delay choices (Fig 5B6). To elucidate the mechanisms that support these dynamics, we performed additional analysis in an exemplar model instance (Fig 7). Again, we concatenated latent factor activity across recordings and choice to perform PCA in this model instance (Fig 7A-7B) and visualized the state projection of latent factor activity onto PC1 and PC2 for latent solution associated with each recording in the testing dataset separately for the immediate (Fig 7A1) and delay (Fig 7A2) choices. This projection was enriched by depicting the speed of trajectories in the latent space (Fig 7B1 and 7B2). In this projection of a single model instance, the differences between immediate and delay choice dynamics were pronounced; immediate choice and delay choice trajectories mostly followed distinct paths in this phase space (Fig 7A1 and 7A2) with distinct profiles in speed of progression (Fig 7B1 and 7B2). Moreover, latent model dynamics were transient: the activity progressed in a counter-clockwise direction from the initial to a final state that marked the end of the trial, speeding up around the time of the choice.

**Figure 7.**
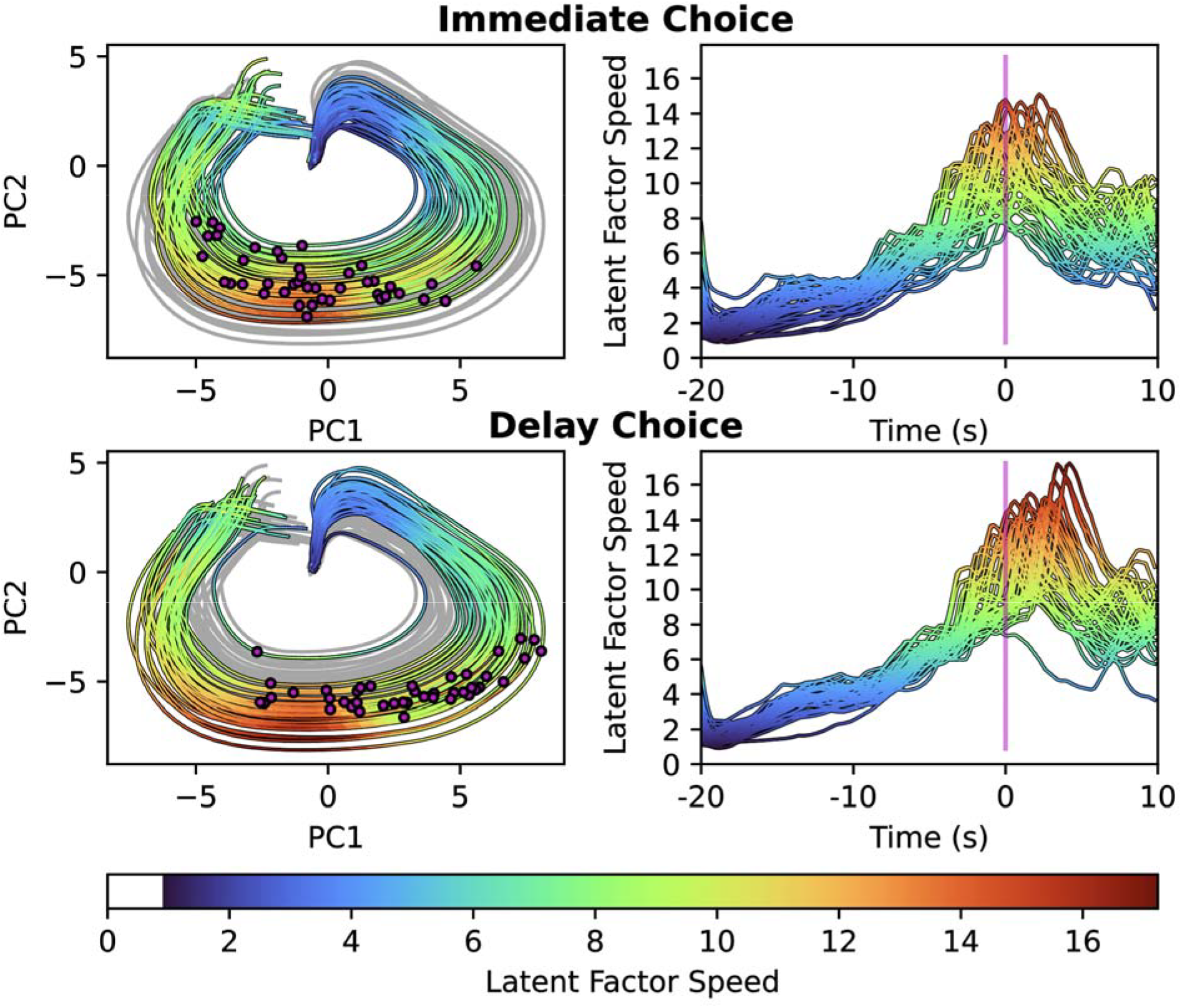
Latent model dynamics differ between immediate and delay choice groups in an exemplar model instance. Projections of latent model activity into the PC1 vs PC2 state space for all immediate choice (A1) and delay choice (A2) solutions for each testing recordings. In each plot, the color-coded trajectories correspond to the choice of interest, and the background grey colored trajectories correspond to the alternate choice. The large purple circular field indicates the neighborhood that contains initial conditions. The purple arrow indicates the direction in which activity evolves. For the trajectory of each recording, we mark the position in PC1-PC2 state space where the choice occurs with a small circular purple marker. Since time is not explicitly handled in part (A), we depict the evolution of the speed of the phase point for each trajectory in the latent state space (B). The vertical purple mark indicates the time of the choice. Color codes the speed in the un-altered latent state space (not in the PC1-PC2 projection of the state space). We computed speed using the difference between successive points in the latent space trajectories divided by the temporal step (200 ms).

### Specialization of latent factor activity determines distinct immediate and delay choice dynamics

To evaluate the role of specific latent factors in neural dynamics and the specialization for immediate and delay choice, we examined latent factors that highly loaded onto the first two principal components in the exemplar model instance (Fig 8). The latent factors with the five highest and five lowest loading coefficients for the first two PCs were selected (Fig 8A1-8A2). Latent factors that were highly loaded onto PC1 exhibited differences in immediate and delay choice trajectories around the time of the choice. Latent factors that were highly loaded onto PC2 exhibited differences in immediate and delay choice trajectories during the interval leading up to the time of the choice and during the reward. These differences are highlighted in the first two principal components (Fig 8C1-8C2) and in the rate at which the high loading latent factors changed (Fig 8D1-8D2). High loading latent factors tended to make choice-and reward-related changes earlier in immediate choices compared to delay choices in PC1 and earlier in delay choices compared to immediate choices in PC2, and notably, both the high-loading latent factors exhibited a peak in speed in the immediate choice shortly after the time of the choice. (Fig 8D1-8D2).

**Figure 8.**
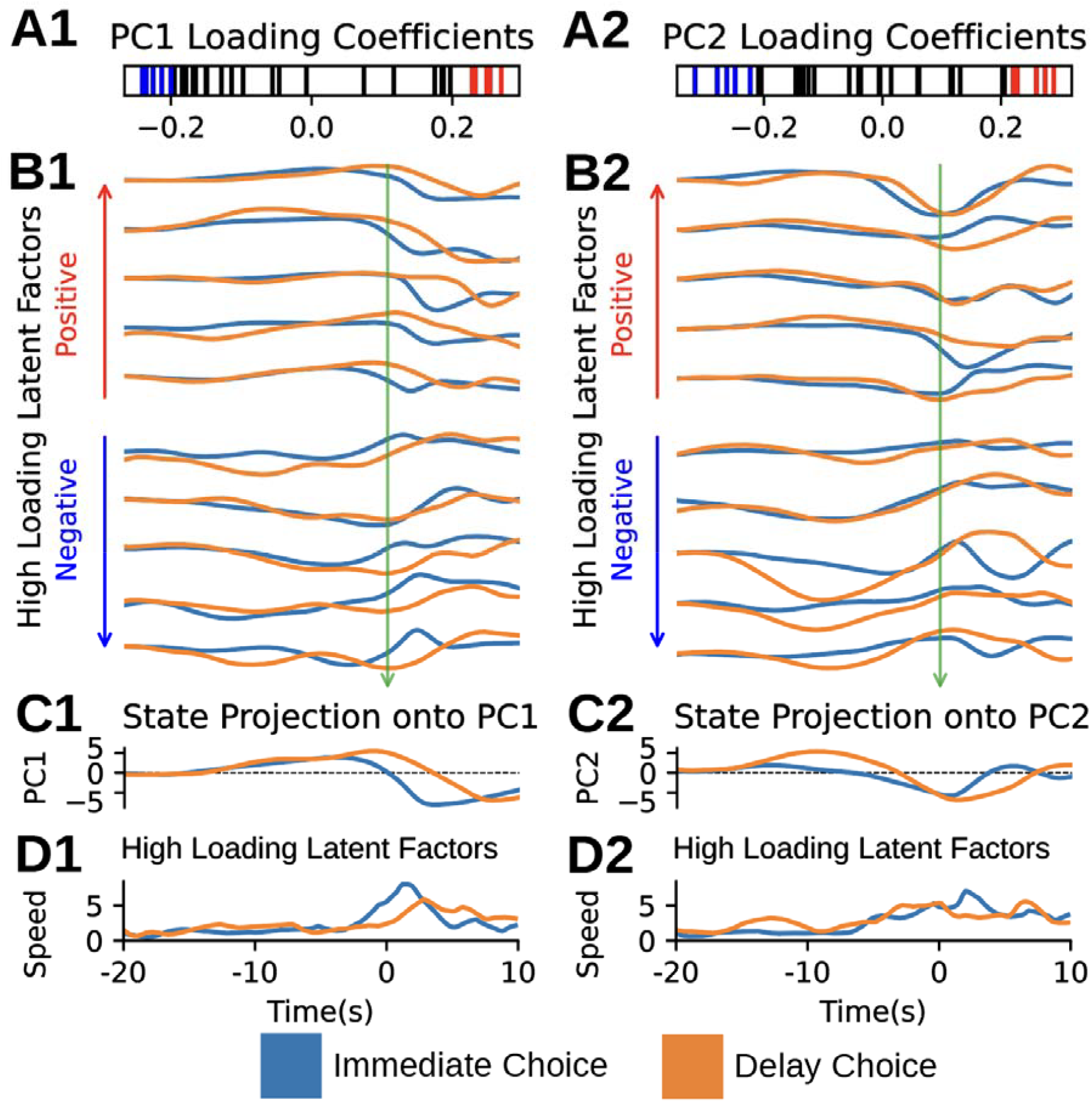
Latent factors differentially participate in the formation of neuronal dynamics for immediate and delay choices. The latent factors with the five highest (red) and five lowest (blue) loading coefficients are highlighted in a loading coefficient plot for the first two PCs (A). High loading latent factors were sorted by their loading coefficients to show groups with similar patterns of activity in immediate and delay choice simulations (B). The first two principal components of latent model activity differentially encoded immediate and delay choice (C). Speed of latent model trajectories in the subspace of the latent space defined by these high loading latent factors (D). We computed speed using the difference between successive points in the high loading latent factors. The vertical green arrow indicates the time of the choice, which is marked as time 0 in the bottom panel (B-D).

To reveal the temporal evolution of the entire latent space, an analysis was performed on the activity of all latent factors, not just the strongest loaders as above. To facilitate visualization of the entire latent model, we sorted its activity by peak absolute value (Fig 9A-9B, right panels). In addition, we depicted the absolute value of PC1 and PC2 loading coefficients for each latent factor (Fig 9A-9B, left panels). The evolution of latent factor activity over time appeared as traveling waves that propagated through the latent units. However, the pattern of loading coefficients differed between choices, indicating that specific latent factors contributed in different phases of latent model activity for different choices. In this way, the PLRNN generated specific and distinct dynamics for immediate and delay choices.

**Figure 9.**
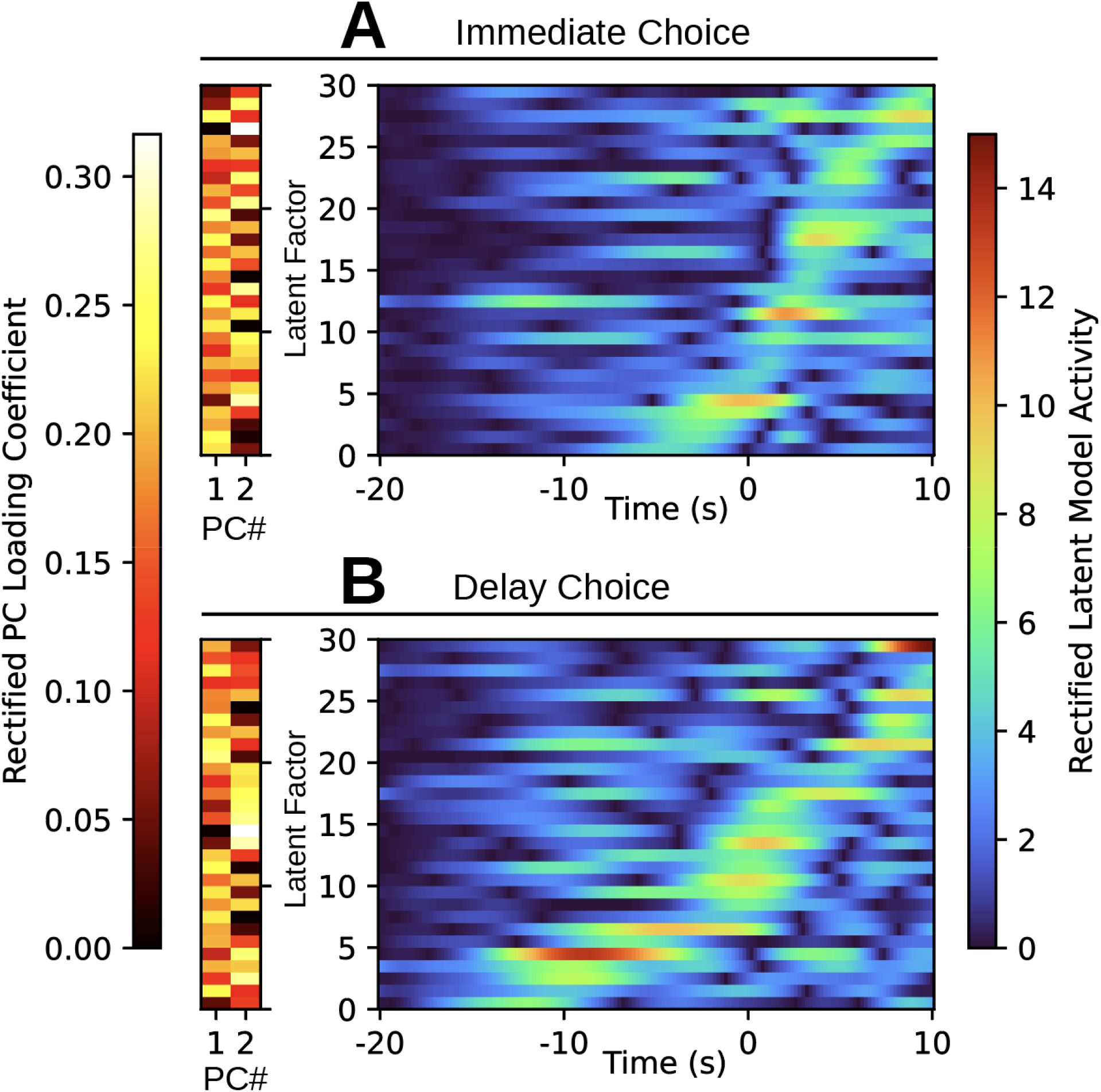
The specialization of latent model activity to generic or choice-specific features of neuronal dynamics is revealed by coding latent factors by their principal component loadings. The absolute value of PC 1 and 2 loading coefficients is coded by the heat scale on the left. The absolute value of the latent model solutions is color coded by the rainbow-colored scale on the right. Latent factors are sorted from top to bottom by the time of peak latent factor activity. The top plot shows immediate choice activity (A), and the bottom plot shows delay choice activity (B). The loading coefficients for PC1 and PC2 are shared across immediate and delay plots, but they appear in a different order according to the order in which latent factors are plotted.

## Discussion

In summary, we have trained a PLRNN model to reproduce neural dynamics for both immediate and delay decisions observed during a delay discounting task. This model extracted common dynamical factors corresponding to the two decisions that generalize across multiple animals. Through cross-validation and the evaluation of overfitting in the observation model, we optimized the size of the latent model. To demonstrate central tendency of model features across instances, we decomposed latent dynamics with principal component analysis and showed how the loading of neurons onto the latent model was similar across model instances.

### The generalization of neural dynamics across datasets from different animals

A primary goal of this work was to show how a PLRNN could be used to generalize neural activity drawn from multiple animals and different recording days into a shared dynamical latent state space. This feat is accomplished here and in other works first by constraining animals to perform the same behavioral tasks, which promotes the recruitment of related motor and cognitive strategies across animals and/or across sessions (Brenner, Weber, et al., 2024; Chen et al., 2021; Herrero-Vidal et al., 2021; Melbaum et al., 2022; Safaie et al., 2023; Schneider et al., 2023). Some of these works utilize a static embedding model, in which the latent representations of neural activity from different experimental sessions are projected onto one another (Chen et al., 2021; Melbaum et al., 2022; Safaie et al., 2023) or formulated in a shared embedding space (Schneider et al., 2023) for comparison. Alternately, data from multiple subjects can be encoded into a dynamical systems model where latent representations from different experimental sessions share a state space (Brenner, Weber, et al., 2024; Herrero-Vidal et al., 2021). In the present work, a PLRNN—a dynamical systems model—was used to jointly model data from multiple sessions drawn from several animals.

Remarkably, instances of our latent model generalized neural activity from disparate experimental sessions into dynamics with apparent rotational structure. Rotational dynamics were first described by Churchland et al. in a reaching task and have since been observed in a variety of motor and cognitive tasks (Aoi et al., 2020; Churchland et al., 2012; Kalidindi et al., 2021; Mante et al., 2013; Nair et al., 2023; Suresh et al., 2020). Recent work has explained rotational dynamics as emerging from traveling waves of activity within a population of neurons (Kuzmina et al., 2024; Lebedev et al., 2019). Using the technique described in Kuzmina et al. (Kuzmina et al., 2024), we characterized the rotational properties of latent model dynamics. Since the present work does not involve a stereotyped motor task, we do not attribute rotational dynamics to the planning and execution of a motor program. Instead, we suggest that rotational dynamics in the latent model state space represent the generalized progression of the animal through different cognitive states of the deliberative decision-making task.

We identified features of latent model activity that differentially encoded choice group: fast activity associated with choice and reward were initiated earlier in immediate choice compared to delay choice (Fig 5B-5C). Through this comparison of neural activity across choice groups, we suggest that cognitive progression across the task is advanced in time during immediate choice compared to delay choice. If this earlier progression of neural activity was only observed after the choice, the advanced temporal progression of immediate choice activity could be attributed to the choice-specific immediate vs delayed encoding of reward. However, our results indicate advanced progression of neural activity for immediate choice tens of seconds before the animal makes a choice, implicating differences in deliberative dynamics.

The cognitive mechanisms underlying deliberation in delay discounting have been modeled as a search process in which the “ease” of finding a solution depends on its location within the topology of a search space (Kurth-Nelson et al., 2012; Linsenbardt et al., 2017). In the search-process framework proposed by Kurth-Nelson et al., outcomes that are less distant into the future are easier to find (Kurth-Nelson et al., 2012). In a drift diffusion model of this framework, the proximity of decision boundaries to the origin was implemented as a function of reward magnitude and delay-time, and initial condition bias was used to model a trial-to-trial decision-making strategy based on previous choices(Linsenbardt et al., 2017). The present work differs from these models in that it does not operate in an explicitly defined decision space; rather, neuronal firing rates were encoded into the state space of a latent model. Despite these differences, the time offset of latent activity between immediate and delay choice observed here could be explained by the relative ease of finding the immediate choice in an abstract deliberative search space.

Since our model generalizes across experimental sessions within animal and between animals and also reproduces neural activity for two different choice groups, we assert that we have achieved a model of the generic computational mechanisms within the ACC that drive the delay discounting task at the level of trial-averaged neural data. Our testing dataset results (Figs 2 and 3) showed that data from unseen recordings can be modeled by computing new recording-specific initial conditions and observation models, and that these unseen recordings were encoded into a shared latent space in such a manner that preserved choice-specific features (Figs 7, 8 and 9). These feats were accomplished using a relatively small portion of the neural recordings available (10 out of 52) to train the latent model, which suggests that many of the features necessary for successful generalization are present in each recording.

### Approaches to dimensionality reduction in dynamical systems models of neural dynamics

Contemporary studies employ several techniques to investigate latent factors that explain features of neural dynamics and animal behavior. In recent work (Nair et al., 2023), neural activity of individual male mice were analyzed with recurrent switching linear dynamical systems models (rSLDS). Similar to PLRNNs, rSLDS dissects the state space of the model into parts with different linear dynamics (Linderman et al., 2017, 2019), but unlike PLRNNs, the dynamics across the borders of linear subregions does not necessarily preserve continuity and the transitions are regulated by a switching variable. Nair et al. (Nair et al., 2023) leveraged rSLDS to identify features of neural dynamics within the ventrolateral subdivision of the ventromedial hypothalamus (VMHvl) and the medial preoptic area (MPOA) that were conserved across models of individual animals. Specifically, behavior was encoded as a line attractor within the VMHvl, and behavior was encoded with rotational dynamics within the MPOA. Pellegrino et al. (Pellegrino et al., 2023) recently described a low-tensor-rank RNN to model neuronal activity across individual trials. Low-rank RNNs address the problems of large parameter spaces and interpretability by imposing low-rank structure on the connectivity matrix (Mastrogiuseppe & Ostojic, 2018). The authors extended the low-rank RNN formulation by implementing the neural network connectivity as a low-rank tensor with parameter vectors that specialize the connectivity matrix to specific trials (Pellegrino et al., 2023). In this way, low-rank tensor RNNs captured the evolution of neural dynamics in macaque motor and premotor cortices over trials as an animal learned a task.

The aforementioned studies employ various ways to confine the model dimensionality and complexity that could also be used for the PLRNN. Although not an inherent property of the model, the rSLDS usually assumes neural activity to be described by just a small number of latent factors (Nair et al., 2023); the low-rank formulation reduces the number of parameters required to construct an RNN connectivity matrix (Mastrogiuseppe & Ostojic, 2018; Pellegrino et al., 2023). While all these assumptions could have easily been implemented in the PLRNN as well (by decomposing or structuring the inter-state connectivity matrix W accordingly), in the present study we decided the model complexity by various statistical criteria, namely finding a model that captures trial-averaged neural activity very faithfully while assuring it is not overfitting the data by comparison with various bootstrap distributions. With this approach, we found an optimal model that describes the dynamics of neural activity from a dataset of 2563 neurons with a set of 30 latent factors.

Essential features of dimensionality reduction are interpretability and generalizability. We adopted additional measures to ensure these features by investigating the consistency of the low dimensional latent space across many instances of our PLRNN model. The intent of our modeling procedure was to capture a minimal representation of neural activity during the decision-making task. If this procedure were successful, then trial-structure and choice-group related activity that exists in the activity of many neurons would be exploited to construct a low-dimensional latent space: not only should our procedure find a low dimensional representation of neural activity, but each instance should capture useful features. Indeed, we found that projections of the latent model activity in PC space were consistent across model instances (Fig 5) and showed how our procedure consistently loaded neurons onto latent states in the observation model (Fig 4).

### Conclusions, Limitations, and Future directions

A major goal of this work was to characterize features of latent dynamics that disambiguate an impulsive from non-impulsive choice. We observed similar dynamics associated with each choice with some nuanced differences. The results of our principal component analysis show differences in phase of principal components between immediate and delay choices (Fig 5B). This feature is emphasized in the PC2-PC1 projection, where the time of the choice happens at a more advanced phase in immediate choices relative to delay choices (Fig 5B and 7A). PC projections also showed how immediate and delay choice neural trajectories traverse different parts of the state space. For example, in an exemplar model instance, the rotational dynamics for delay choice trajectories were more likely to traverse farther out into the PC2-PC1 projection (Fig 7A) and accelerate later in the rotational phase (Fig 7B) than immediate choice trajectories. These differences were also apparent in individual latent factors; PC1- and PC2-high loading latent factors consistently and differentially encoded immediate and delay choice (Fig 8) and were active in different phases of neural activity progression between immediate and delay choice (Fig 9). A key takeaway from this work is that our latent factor analysis indicated choice-related neural activation earlier in the task interval for immediate choices than for late choices (Fig 5B and Fig 8D); these results implicate earlier fast dynamics in impulsive choice compared to non-impulsive choice.

It is important to note that in the present context we harvested the PLRNN more as a sophisticated form of nonlinear factor analysis that captures essential trial-average activity profiles, rather than performing dynamical systems reconstruction as defined in Durstewitz et al. (Durstewitz et al., 2023). Dynamical systems reconstruction that account for single trial evolution of the dynamics using recent algorithms like generalized or multimodal teacher forcing (Brenner, Hess, et al., 2024; Hess et al., 2023), or methods for directly integrating dynamics across many different experimental subjects (Brenner, Hess, et al., 2024; Brenner, Weber, et al., 2024) is a necessary step in clearly linking the neural dynamics that underlie behavior.

## Methods

### PLRNN definition

The PLRNN model consisted of two modules: a latent model of neural dynamics and an observation model that mapped the latent variables onto the observed spike trains. The latent model (with latent states ***z***_***t***_) was shared across all recordings in the training and testing datasets, and unique instances of the observation model (with observed variables ***x***_***t***_) transformed the latent model dynamics to reproduce the single unit firing rates during immediate and delay decisions for each recording:

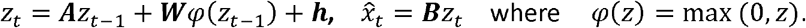

The latent state (***z***_*t*_) is an M-dimensional vector, and the number of elements was a hyperparameter *M* that we varied from 3 to 1000 variables. The observation model represented the firing rates of single units. ***A*** is a *M×M* diagonal matrix of autoregressive weights; ***W*** a *M×M* off-diagonal matrix containing the neural network connectivity weights; ***h*** a vector of length *M* containing constants that determined the excitability. ***B*** is a matrix of regression weights connecting the latent states to the observed. During training, ***B*** was partitioned by recording. Each such partition was *NΩM* according to the number of neurons in that recording (*N*). The latent model initial conditions (*μ*_0_) were similarly trained as a parameter. The initial condition was specific for each recording and separate for the immediate vs. delay conditions. The Jacobian of the latent model was computed as *J*_*t*_ =*A* + *wD*_Ω (t-1)_ such that *D*_Ω (t)_= diag(*d*_Ω(t)_) and *d*_Ω(t)_=(*d*_1_,*d*_2_ ,…*d*_m_,) . The vector *d*Ω_(t)_ indicated *dm* (z_*m*_,_*t*_)= 1 where z _*m,t*_ >0 and zero otherwise. In this way, the piecewise linear activations yield 2^*m*^ linear subregions within the state space z.

### Training of PLRNNs

We used backpropagation through time to fit the parameters contained in ***A, W, h, B***, and ***μ***_0_. The loss was computed as the average of the mean absolute error of predicted firing rates over each choice group for each neuron. Parameters were optimized using the Adam optimizer. Several of our hyperparameters were determined following some rounds of preliminary optimization. The learning rate was set to 1e-3 and other parameters of the optimizer took the default values as provided by the Optax software package. In each optimization epoch, the **B** matrices for each recording were concatenated and vectorized to perform regularization with an L1 and an L2 penalty. These quantities were weighted (in this case evenly each with the coefficient 0.5), were multiplied by a regularization coefficient (1e-5), and were added to the loss. We employed update clipping with a clipping size of 0.2. The fitting of parameters was performed over 7000 epochs for the training dataset and 3000 epochs for the testing dataset. Ten recordings were used for the training dataset and forty-two recordings were used for the testing dataset. Since the initial condition (***μ***_0_) and observation model (B) parameters were specialized by the recording to specific neurons, it was necessary to fit these parameters “from scratch” for the testing dataset. In this case, the latent model parameters ***A, W***, and ***h*** were “frozen” while ***μ***_0_ and ***B*** were fit for each recording in the testing dataset.

### Analysis of PLRNN

To evaluate overfitting in the observation model, we scrambled the off-diagonal elements of the matrix ***W*** and refit the initial condition and observation model using the same procedure as used for the testing dataset. To further evaluate overfitting in the observation model, we randomly shuffled the latent model solution in chunks of 5 and 15 time steps. These surrogate latent model solutions were used to fit an observation model (but not an initial condition) with optimization hyperparameters drawn from the optimization routine for the testing dataset as described above.

In Figure 3A, we depict the results for bootstrap tests between two distributions for overlapping IQR. In each bootstrap sample, we computed the first quartile (Q1) for the testing distribution and the third quartile (Q3) for the designated comparison distribution (Scramble-W, Shuffle 5, or Shuffle 15). We counted how many times over all bootstrap samples the testing distribution Q1 was less than the comparison distribution Q3. To derive a p-value, this count was divided by the number of bootstrap samples. We used 10000 bootstrap samples.

The dynamics of the neural activity and the latent model were analyzed with principal component analysis (PCA) using the singular value decomposition method from Numpy’s linear algebra module (numpy.linalg.svd). PCA was performed on neural activity time series that were concatenated across choice conditions and then z-scored. PCA was performed on latent model time series that were concatenated across testing recordings and choice conditions and then z-scored. For Figure 5, we found that the sign of the loading coefficients for similar principal components in different model instances were flipped compared to one another. We used an agglomerative clustering routine to pick out 2 clusters among these principal components and multiplied the components in one cluster by -1 to align them in the same direction as the neural activity PCs in Fig 5A for visualization.

To evaluate the stability of our training routine across model instances, we used consensus clustering to analyze the loadings of individual neurons in the observation model (the matrix ***B***; 4B-4C). This matrix had 2563 rows (corresponding to each neuron) each containing a number of features corresponding to size of the latent model (2563xM), and clustering was performed in row space (i.e., across neuronal latent loading vectors). For each model instance, k-means was used to group neurons into 20 clusters, and we counted how many times did each neuron share a cluster with each other neuron over all 100 model instances. By normalizing this count, we produced the clustering likelihood estimation. To visualize these estimations, we reduced these estimations to histograms with 101 bins from 0 to 100 percent likelihood. The clustering likelihood estimates were tested against one hundred independently generated null models. To produce a` null model, we shuffled the cluster labels and recomputed the clustering likelihood estimation. We tested observations against the null models with the two-sample Kolmogorov-Smirnov test in the statistics module of Scipy (scipy.stats.ks_2samp) using the ‘exact’ method argument. In figure 5A, we first reduced the 2563xM matrix to 2563×2 with tSNE before grouping neurons into 10 clusters; we used the tSNE routine from scikit-learn (sklearn.manifold.TSNE) with the arguments n_components=2, metric=“cosine”, perplexity=30. This panel is intended as a visual aide to help understand the procedure described above.

For figure 7, we selected an exemplar model based on visual appeal. We examined the top 10 models based on testing group MAE and selected the 4^th^ best model for visualization. This model instance was also used in figures 8 and 9.

To quantify rotational dynamics in the latent model state space, we computed the gyration number as described in Kuzmina et al. (Kuzmina et al., 2024). Data were first mean-centered across sessions at each time point and concatenated across sessions and conditions. The gyration number was then derived from the eigenvalues (*λ*_*i*_) of the neuron differential covariance matrix: 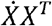 . The gyration number provided the coordinates in the Gyration Plane, which were computed from the two largest eigenvalues as

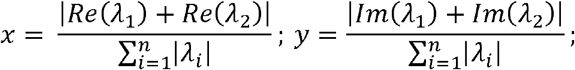

where *x* and *y* respectively are the Decay Axis and Rotation Axis in the Gyration plane. This analysis was performed using source code directly adapted from the repository (https://github.com/NevVerVer/neural-dynamics-gyration) provided by Kuzmina et al. (Kuzmina et al., 2024).

In all figures that include box-and-whisker plots, we have excluded outliers from diagrams.

### Software

RNN simulations and optimizations were implemented using the Flax, Optax, and JAX python packages. Data and model analysis routines also included Numpy, Scipy, Scikit-learn, and Pandas python packages. Figures were produced using Matplotlib and edited in Adobe Illustrator. The data and software used to produce this work are available at https://github.com/whbdupree/DelayDiscountingPLRNN.

## Acknowledgements

We thank Seth Weir and Jacob Worrell for their contributions to the source code used in this project. This work was supported by grants to CCL from NIH (AA029970, AA029409), and DD was supported by grants from the German Science Foundation (DFG Du 354/18-1, Du 354/15-1, and Du 354/14-1 within the FOR-5159). This research was supported in part by Lilly Endowment, Inc., through its support for the Indiana University Pervasive Technology Institute.

---

**Figure S1.**
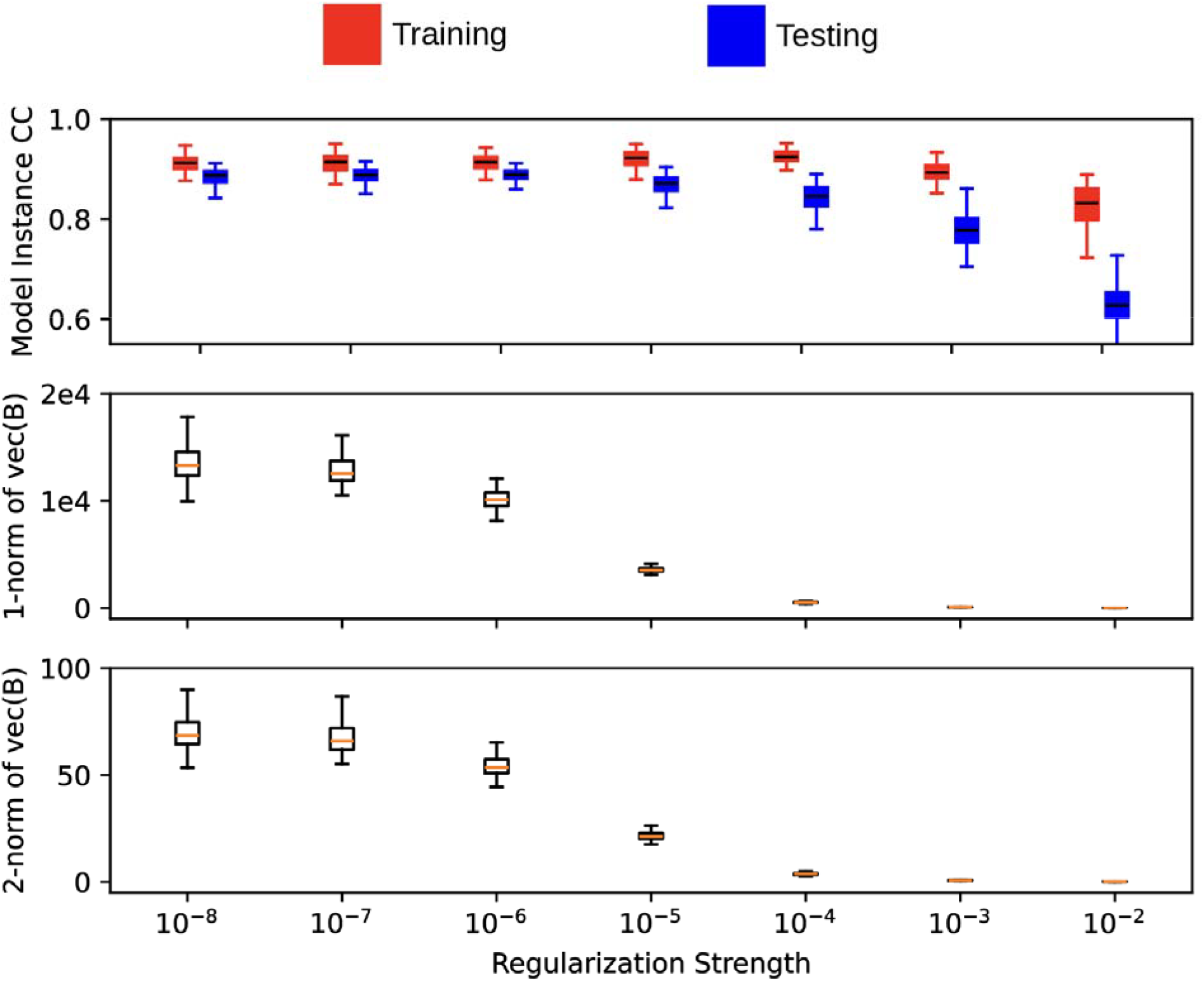
Selection of regularization penalty. We selected its value using the elbow criterion to balance model performance, the 1-norm of vec(**B**), and the 2-norm of vec(**B**). The value used for all other results is 10^-5^. All distributions show cross validation across 100 model instances.

**Figure S2.**
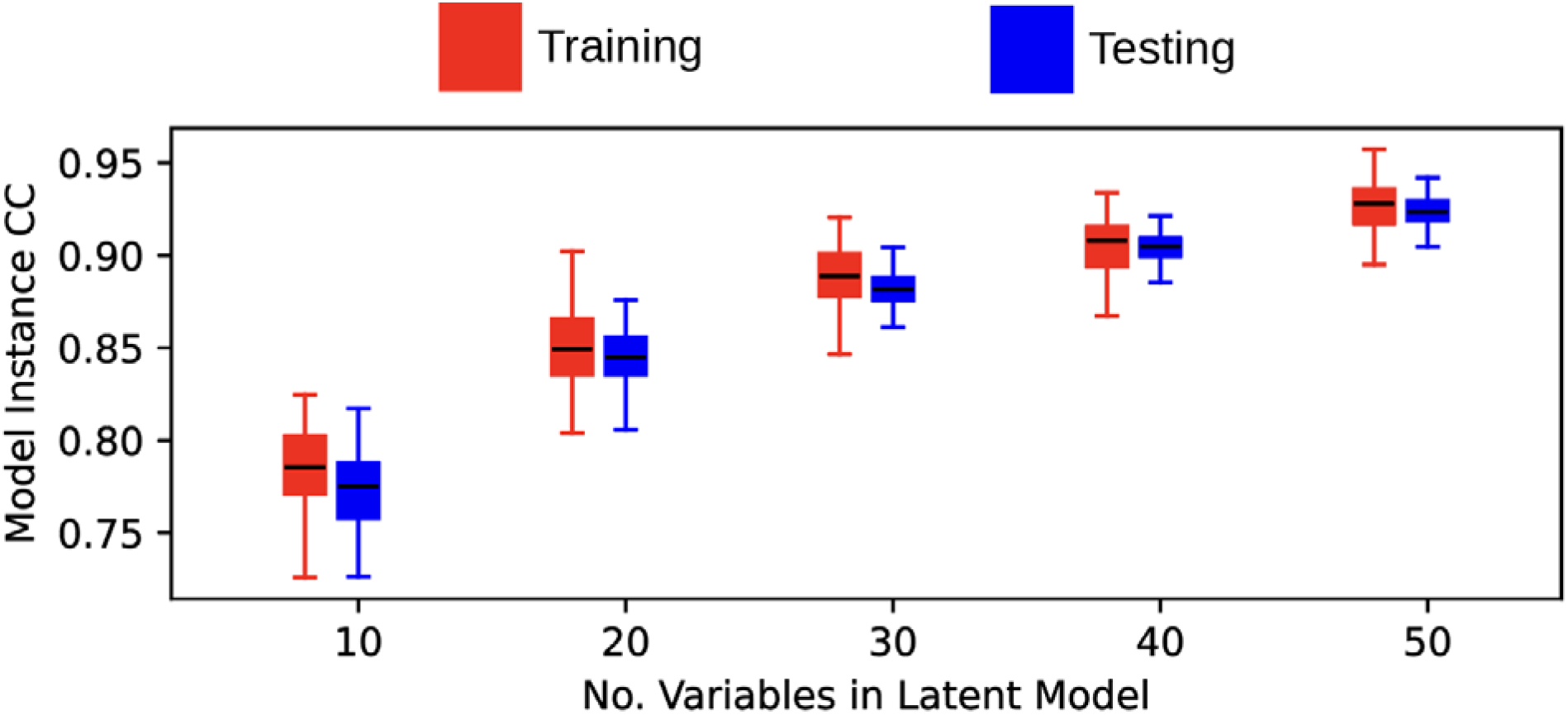
Linear latent factor model performance for different model sizes. Model instance correlation coefficient (CC) was determined by the correlation coefficient of the generated and observed spike trains and then averaged across neurons and across immediate and delay choices to produce a single value for each model instance. The red distributions correspond to training dataset performance, and the blue distributions correspond to testing dataset performance.

**Figure S3.**
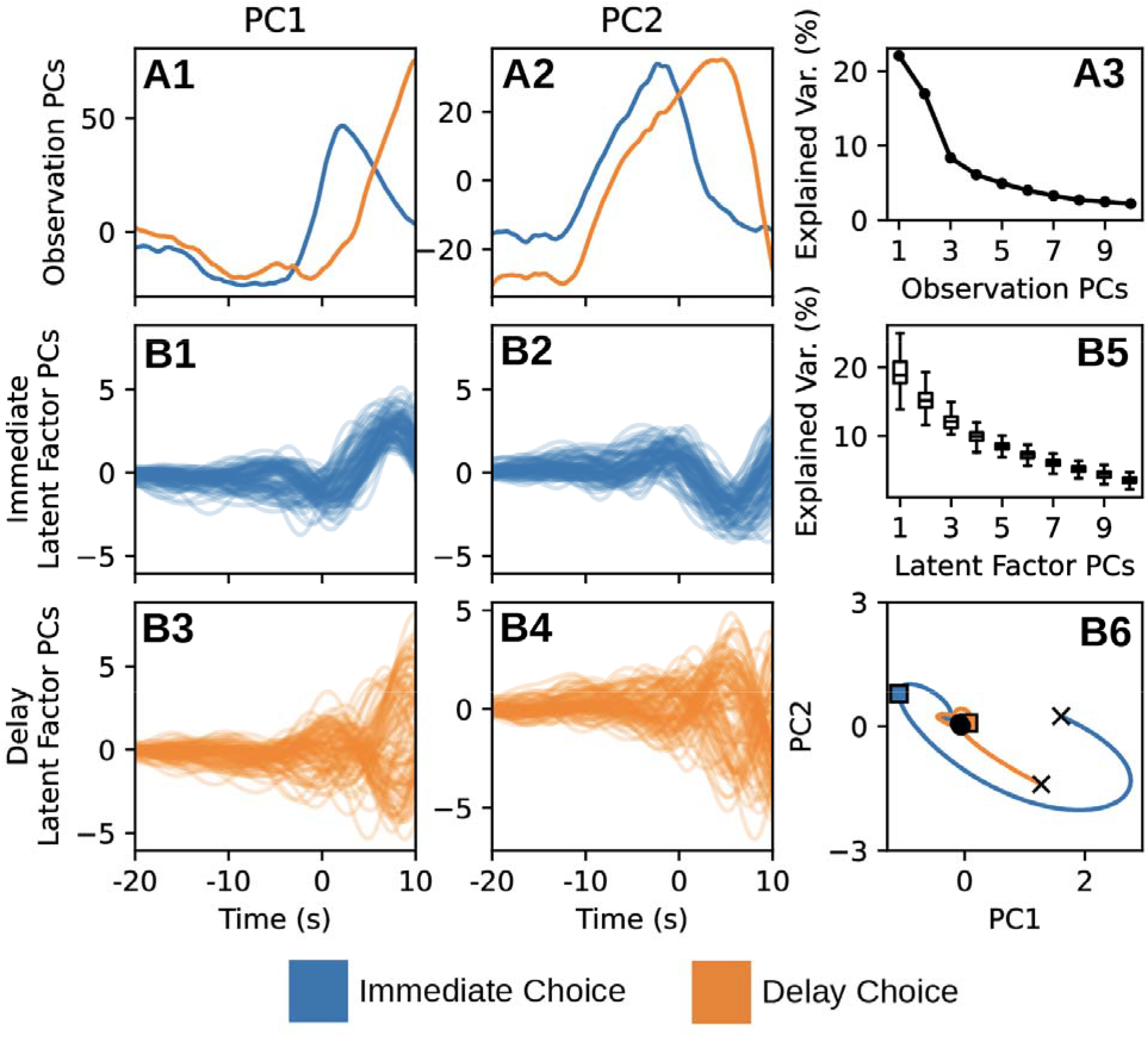
Latent model dynamics for the linear latent factor model (LRNN) across model instances. Blue traces correspond to immediate choice activity, and orange traces correspond to delay choice activity. (A) Top row traces indicate results of PCA on neuronal depicting the PC1 (A1) and PC2 (A2) projections for immediate choice and delay choice. The combined explained variance of neuronal activity for the first two PCs was less than 50% (A3). PCA was performed separately on latent model activity for each model instance (B). Each colored line corresponds to the mean of the projections onto PC1 or PC2 for each of the 42 testing recordings within a model instance (B1-B4; one trace for each of 100 model instances). The distribution of explained variance for the top 10 PCs for all model instances is visualized (B5). The mean of the traces from B1-B4 are adopted as generalized neuronal dynamics in the PC1-PC2 plot (B6). Within this projection the progression of the state through time is marked: circles which correspond to the initial state; filled squares which indicate the time of the choice (time = 0); and Xs, which indicate the final state.

## Works Cited

Aoi, M. C., Mante, V., & Pillow, J. W. (2020). Prefrontal cortex exhibits multidimensional dynamic encoding during decision-making. Nature Neuroscience, 23(11), 1410–1420. 10.1038/s41593-020-0696-5

Balaguer-Ballester, E., Lapish, C. C., Seamans, J. K., & Durstewitz, D. (2011). Attracting dynamics of frontal cortex ensembles during memory-guided decision-making. PLoS Computational Biology, 7(5), e1002057. 10.1371/journal.pcbi.1002057

Brenner, M., Hess, F., Koppe, G., & Durstewitz, D. (2024). Integrating Multimodal Data for Joint Generative Modeling of Complex Dynamics. Proceedings of the 41st International Conference on Machine Learning, 4482–4516. https://proceedings.mlr.press/v235/brenner24a.html

Brenner, M., Weber, E., Koppe, G., & Durstewitz, D. (2024). Learning Interpretable Hierarchical Dynamical Systems Models from Time Series Data (Version 2). arXiv. 10.48550/ARXIV.2410.04814

Brody, C. D., & Hanks, T. D. (2016). Neural underpinnings of the evidence accumulator. Current Opinion in Neurobiology, 37, 149–157. 10.1016/j.conb.2016.01.003

Carr, D. B., & Sesack, S. R. (2000). Projections from the rat prefrontal cortex to the ventral tegmental area: Target specificity in the synaptic associations with mesoaccumbens and mesocortical neurons. The Journal of Neuroscience: The Official Journal of the Society for Neuroscience, 20(10), 3864–3873. 10.1523/JNEUROSCI.20-10-03864.2000

Chen, H.-T., Manning, J. R., & van der Meer, M. A. A. (2021). Between-subject prediction reveals a shared representational geometry in the rodent hippocampus. Current Biology: CB, 31(19), 4293–4304.e5. 10.1016/j.cub.2021.07.061

Chow, T. W. S., & Li, X.-D. (2000). Modeling of continuous time dynamical systems with input by recurrent neural networks. IEEE Transactions on Circuits and Systems I: Fundamental Theory and Applications, 47(4), 575–578. 10.1109/81.841860

Churchland, M. M., Cunningham, J. P., Kaufman, M. T., Foster, J. D., Nuyujukian, P., Ryu, S. I., & Shenoy, K. V. (2012). Neural population dynamics during reaching. Nature, 487(7405), 51–56. 10.1038/nature11129

Churchwell, J. C., Morris, A. M., Heurtelou, N. M., & Kesner, R. P. (2009). Interactions Between the Prefrontal Cortex and Amygdala During Delay Discounting and Reversal. Behavioral Neuroscience, 123(6), 1185–1196. 10.1037/a0017734

Cunningham, J. P., & Yu, B. M. (2014). Dimensionality reduction for large-scale neural recordings. Nature Neuroscience, 17(11), 1500–1509. 10.1038/nn.3776

Dalley, J. W., Everitt, B. J., & Robbins, T. W. (2011). Impulsivity, compulsivity, and top-down cognitive control. Neuron, 69(4), 680–694. 10.1016/j.neuron.2011.01.020

Durstewitz, D. (2017). A state space approach for piecewise-linear recurrent neural networks for identifying computational dynamics from neural measurements. PLoS Computational Biology, 13(6), e1005542. 10.1371/journal.pcbi.1005542

Durstewitz, D., Koppe, G., & Thurm, M. I. (2023). Reconstructing computational system dynamics from neural data with recurrent neural networks. Nature Reviews Neuroscience, 24(11), 693–710. 10.1038/s41583-023-00740-7

Euston, D. R., Gruber, A. J., & McNaughton, B. L. (2012). The role of medial prefrontal cortex in memory and decision making. Neuron, 76(6), 1057–1070. 10.1016/j.neuron.2012.12.002

Floresco, S. B., Seamans, J. K., & Phillips, A. G. (1997). Selective roles for hippocampal, prefrontal cortical, and ventral striatal circuits in radial-arm maze tasks with or without a delay. The Journal of Neuroscience: The Official Journal of the Society for Neuroscience, 17(5), 1880–1890. 10.1523/JNEUROSCI.17-05-01880.1997

Funahashi, K., & Nakamura, Y. (1993). Approximation of dynamical systems by continuous time recurrent neural networks. Neural Networks, 6(6), 801–806. 10.1016/S0893-6080(05)80125-X

Gallego, J. A., Perich, M. G., Miller, L. E., & Solla, S. A. (2017). Neural Manifolds for the Control of Movement. Neuron, 94(5), 978–984. 10.1016/j.neuron.2017.05.025

Gisquet-Verrier, P., & Delatour, B. (2006). The role of the rat prelimbic/infralimbic cortex in working memory: Not involved in the short-term maintenance but in monitoring and processing functions. Neuroscience, 141(2), 585–596. 10.1016/j.neuroscience.2006.04.009

Gold, J. I., & Shadlen, M. N. (2007). The neural basis of decision making. Annual Review of Neuroscience, 30, 535–574. 10.1146/annurev.neuro.29.051605.113038

Goto, Y., & Grace, A. A. (2008). Dopamine modulation of hippocampal-prefrontal cortical interaction drives memory-guided behavior. Cerebral Cortex (New York, N.Y.: 1991), 18(6), 1407–1414. 10.1093/cercor/bhm172

Hanks, T. D., & Summerfield, C. (2017). Perceptual Decision Making in Rodents, Monkeys, and Humans. Neuron, 93(1), 15–31. 10.1016/j.neuron.2016.12.003

Herrero-Vidal, P., Rinberg, D., & Savin, C. (2021). Across-animal odor decoding by probabilistic manifold alignment. Advances in Neural Information Processing Systems, 34, 20360–20372. https://proceedings.neurips.cc/paper_files/paper/2021/hash/aad64398a969ec3186800d412fa7ab31-Abstract.html

Hess, F., Monfared, Z., Brenner, M., & Durstewitz, D. (2023). Generalized Teacher Forcing for Learning Chaotic Dynamics. Proceedings of the 40th International Conference on Machine Learning, 13017–13049. https://proceedings.mlr.press/v202/hess23a.html

Hoover, W. B., & Vertes, R. P. (2007). Anatomical analysis of afferent projections to the medial prefrontal cortex in the rat. Brain Structure & Function, 212(2), 149–179. 10.1007/s00429-007-0150-4

Kalidindi, H. T., Cross, K. P., Lillicrap, T. P., Omrani, M., Falotico, E., Sabes, P. N., & Scott, S. H. (2021). Rotational dynamics in motor cortex are consistent with a feedback controller. eLife, 10, e67256. 10.7554/eLife.67256

Keshtkaran, M. R., Sedler, A. R., Chowdhury, R. H., Tandon, R., Basrai, D., Nguyen, S. L., Sohn, H., Jazayeri, M., Miller, L. E., & Pandarinath, C. (2022). A large-scale neural network training framework for generalized estimation of single-trial population dynamics. Nature Methods, 19(12), 1572–1577. 10.1038/s41592-022-01675-0

Kimura, M., & Nakano, R. (1998). Learning dynamical systems by recurrent neural networks from orbits. Neural Networks, 11(9), 1589–1599. 10.1016/S0893-6080(98)00098-7

Koppe, G., Toutounji, H., Kirsch, P., Lis, S., & Durstewitz, D. (2019). Identifying nonlinear dynamical systems via generative recurrent neural networks with applications to fMRI. PLOS Computational Biology, 15(8), e1007263. 10.1371/journal.pcbi.1007263

Kurikawa, T., Haga, T., Handa, T., Harukuni, R., & Fukai, T. (2018). Neuronal stability in medial frontal cortex sets individual variability in decision-making. Nature Neuroscience, 21(12), 1764–1773. 10.1038/s41593-018-0263-5

Kurth-Nelson, Z., Bickel, W., & Redish, A. D. (2012). A theoretical account of cognitive effects in delay discounting. The European Journal of Neuroscience, 35(7), 1052–1064. 10.1111/j.1460-9568.2012.08058.x

Kuzmina, E., Kriukov, D., & Lebedev, M. (2024). Neuronal travelling waves explain rotational dynamics in experimental datasets and modelling. Scientific Reports, 14(1), 3566. 10.1038/s41598-024-53907-2

Lapish, C. C., Balaguer-Ballester, E., Seamans, J. K., Phillips, A. G., & Durstewitz, D. (2015). Amphetamine Exerts Dose-Dependent Changes in Prefrontal Cortex Attractor Dynamics during Working Memory. The Journal of Neuroscience: The Official Journal of the Society for Neuroscience, 35(28), 10172–10187. 10.1523/JNEUROSCI.2421-14.2015

Lapish, C. C., Durstewitz, D., Chandler, L. J., & Seamans, J. K. (2008). Successful choice behavior is associated with distinct and coherent network states in anterior cingulate cortex. Proceedings of the National Academy of Sciences of the United States of America, 105(33), 11963–11968. 10.1073/pnas.0804045105

Lebedev, M. A., Ossadtchi, A., Mill, N. A., Urpí, N. A., Cervera, M. R., & Nicolelis, M. A. L. (2019). Analysis of neuronal ensemble activity reveals the pitfalls and shortcomings of rotation dynamics. Scientific Reports, 9(1), 18978. 10.1038/s41598-019-54760-4

Lévesque, M., & Parent, A. (1998). Axonal arborization of corticostriatal and corticothalamic fibers arising from prelimbic cortex in the rat. Cerebral Cortex (New York, N.Y.: 1991), 8(7), 602–613. 10.1093/cercor/8.7.602

Linderman, S., Johnson, M., Miller, A., Adams, R., Blei, D., & Paninski, L. (2017). Bayesian Learning and Inference in Recurrent Switching Linear Dynamical Systems. Proceedings of the 20th International Conference on Artificial Intelligence and Statistics, 914–922. https://proceedings.mlr.press/v54/linderman17a.html

Linderman, S., Nichols, A., Blei, D., Zimmer, M., & Paninski, L. (2019). Hierarchical recurrent state space models reveal discrete and continuous dynamics of neural activity in C. elegans (p. 621540). bioRxiv. 10.1101/621540

Linsenbardt, D. N., Smoker, M. P., Janetsian-Fritz, S. S., & Lapish, C. C. (2017). Impulsivity in rodents with a genetic predisposition for excessive alcohol consumption is associated with a lack of a prospective strategy. Cognitive, Affective, & Behavioral Neuroscience, 17(2), 235–251. 10.3758/s13415-016-0475-7

Loos, M., Pattij, T., Janssen, M. C. W., Counotte, D. S., Schoffelmeer, A. N. M., Smit, A. B., Spijker, S., & van Gaalen, M. M. (2010). Dopamine receptor D1/D5 gene expression in the medial prefrontal cortex predicts impulsive choice in rats. Cerebral Cortex (New York, N.Y.: 1991), 20(5), 1064–1070. 10.1093/cercor/bhp167

Macke, J. H., Buesing, L., Cunningham, J. P., Yu, B. M., Shenoy, K. V., & Sahani, M. (2011). Empirical models of spiking in neural populations. Advances in Neural Information Processing Systems, 24. https://papers.nips.cc/paper_files/paper/2011/hash/7143d7fbadfa4693b9eec507d9d37443-Abstract.html

Mante, V., Sussillo, D., Shenoy, K. V., & Newsome, W. T. (2013). Context-dependent computation by recurrent dynamics in prefrontal cortex. Nature, 503(7474), 78–84. 10.1038/nature12742

Mastrogiuseppe, F., & Ostojic, S. (2018). Linking Connectivity, Dynamics, and Computations in Low-Rank Recurrent Neural Networks. Neuron, 99(3), 609–623.e29. 10.1016/j.neuron.2018.07.003

McLaughlin, A. E., & Redish, A. D. (2023). Optogenetic disruption of the prelimbic cortex alters long-term decision strategy but not valuation on a spatial delay discounting task. Neurobiology of Learning and Memory, 200, 107734. 10.1016/j.nlm.2023.107734

Melbaum, S., Russo, E., Eriksson, D., Schneider, A., Durstewitz, D., Brox, T., & Diester, I. (2022). Conserved structures of neural activity in sensorimotor cortex of freely moving rats allow cross-subject decoding. Nature Communications, 13(1), Article 1. 10.1038/s41467-022-35115-6

Miller, P. (2016). Dynamical systems, attractors, and neural circuits. F1000Research, 5, F1000 Faculty Rev-992. 10.12688/f1000research.7698.1

Myroshnychenko, M., Seamans, J. K., Phillips, A. G., & Lapish, C. C. (2017). Temporal Dynamics of Hippocampal and Medial Prefrontal Cortex Interactions During the Delay Period of a Working Memory-Guided Foraging Task. Cerebral Cortex (New York, N.Y.: 1991), 27(11), 5331–5342. 10.1093/cercor/bhx184

Nair, A., Karigo, T., Yang, B., Ganguli, S., Schnitzer, M. J., Linderman, S. W., Anderson, D. J., & Kennedy, A. (2023). An approximate line attractor in the hypothalamus encodes an aggressive state. Cell, 186(1), 178–193.e15. 10.1016/j.cell.2022.11.027

Nonnenmacher, M., Turaga, S. C., & Macke, J. H. (2017). Extracting low-dimensional dynamics from multiple large-scale neural population recordings by learning to predict correlations. Advances in Neural Information Processing Systems, 30. https://papers.nips.cc/paper_files/paper/2017/hash/c7558e9d1f956b016d1fdba7ea132378-Abstract.html

Pals, M., Sağtekin, A. E., Pei, F., Gloeckler, M., & Macke, J. H. (2024). Inferring stochastic low-rank recurrent neural networks from neural data (arXiv:2406.16749; Version 1). arXiv. 10.48550/arXiv.2406.16749

Pandarinath, C., O’Shea, D. J., Collins, J., Jozefowicz, R., Stavisky, S. D., Kao, J. C., Trautmann, E. M., Kaufman, M. T., Ryu, S. I., Hochberg, L. R., Henderson, J. M., Shenoy, K. V., Abbott, L. F., & Sussillo, D. (2018). Inferring single-trial neural population dynamics using sequential autoencoders. Nature Methods, 15(10), 805–815. 10.1038/s41592-018-0109-9

Patel, A. N., Sedler, A. R., Huang, J., Pandarinath, C., & Gilja, V. (2023). High-performance neural population dynamics modeling enabled by scalable computational infrastructure. Journal of Open Source Software, 8(83), 5023. 10.21105/joss.05023

Pellegrino, A., Cayco Gajic, N. A., & Chadwick, A. (2023). Low Tensor Rank Learning of Neural Dynamics. Advances in Neural Information Processing Systems, 36, 11674–11702.

Perich, M. G., Arlt, C., Soares, S., Young, M. E., Mosher, C. P., Minxha, J., Carter, E., Rutishauser, U., Rudebeck, P. H., Harvey, C. D., & Rajan, K. (2021). Inferring brain-wide interactions using data-constrained recurrent neural network models (p. 2020.12.18.423348). bioRxiv. 10.1101/2020.12.18.423348

Rabinovich, M. I., & Varona, P. (2011). Robust transient dynamics and brain functions. Frontiers in Computational Neuroscience, 5, 24. 10.3389/fncom.2011.00024

Safaie, M., Chang, J. C., Park, J., Miller, L. E., Dudman, J. T., Perich, M. G., & Gallego, J. A. (2023). Preserved neural dynamics across animals performing similar behaviour. Nature, 623(7988), 765–771. 10.1038/s41586-023-06714-0

Schneider, S., Lee, J. H., & Mathis, M. W. (2023). Learnable latent embeddings for joint behavioural and neural analysis. Nature, 617(7960), 360–368. 10.1038/s41586-023-06031-6

Seamans, J. K., White, S., Morningstar, M., Emberly, E., Linsenbardt, D., Ma, B., Czachowski, C. L., & Lapish, C. C. (2024). Neural basis of cognitive control signals in anterior cingulate cortex during delay discounting (p. 2024.06.07.597894). bioRxiv. 10.1101/2024.06.07.597894

Sedler, A. R., & Pandarinath, C. (2023). lfads-torch: A modular and extensible implementation of latent factor analysis via dynamical systems (arXiv:2309.01230). arXiv. 10.48550/arXiv.2309.01230

Suresh, A. K., Goodman, J. M., Okorokova, E. V., Kaufman, M., Hatsopoulos, N. G., & Bensmaia, S. J. (2020). Neural population dynamics in motor cortex are different for reach and grasp. eLife, 9, e58848. 10.7554/eLife.58848

Sussillo, D., Churchland, M. M., Kaufman, M. T., & Shenoy, K. V. (2015). A neural network that finds a naturalistic solution for the production of muscle activity. Nature Neuroscience, 18(7), 1025–1033. 10.1038/nn.4042

Timme, N. M., Ma, B., Linsenbardt, D., Cornwell, E., Galbari, T., & Lapish, C. C. (2022). Compulsive alcohol drinking in rodents is associated with altered representations of behavioral control and seeking in dorsal medial prefrontal cortex. Nature Communications, 13, 3990. 10.1038/s41467-022-31731-4

Valente, A., Pillow, J. W., & Ostojic, S. (2022). Extracting computational mechanisms from neural data using low-rank RNNs. Advances in Neural Information Processing Systems, 35, 24072–24086.

van Heukelum, S., Mars, R. B., Guthrie, M., Buitelaar, J. K., Beckmann, C. F., Tiesinga, P. H. E., Vogt, B. A., Glennon, J. C., & Havenith, M. N. (2020). Where is Cingulate Cortex? A Cross-Species View. Trends in Neurosciences, 43(5), 285–299. 10.1016/j.tins.2020.03.007

White, S. M., Morningstar, M. D., De Falco, E., Linsenbardt, D. N., Ma, B., Parks, M. A., Czachowski, C. L., & Lapish, C. C. (2023). Flexible coding schemes in dorsomedial prefrontal cortex underlie decision-making during delay discounting. bioRxiv, 2023.06.15.545101. 10.1101/2023.06.15.545101

Wong, K.-F., & Wang, X.-J. (2006). A recurrent network mechanism of time integration in perceptual decisions. The Journal of Neuroscience: The Official Journal of the Society for Neuroscience, 26(4), 1314–1328. 10.1523/JNEUROSCI.3733-05.2006

Zhao, Y., & Park, I. M. (2016). Interpretable Nonlinear Dynamic Modeling of Neural Trajectories. Advances in Neural Information Processing Systems, 29. https://proceedings.neurips.cc/paper/2016/hash/b2531e7bb29bf22e1daae486fae3417a-Abstract.html

